# Diverse gene ancestries reveal multiple microbial associations during eukaryogenesis

**DOI:** 10.1101/2024.10.14.618062

**Authors:** Moisès Bernabeu, Saioa Manzano-Morales, Marina Marcet-Houben, Toni Gabaldón

**Author notes:** These authors contributed equally and their names are listed in alphabetical order (Moisès Bernabeu, Saioa Manzano-Morales, Marina Marcet-Houben).

## Abstract

The origin of eukaryotes remains a central enigma in biology^1^. Ongoing debates agree on the pivotal role of a symbiosis between an alphaproteobacterium and an Asgard archaeon^2,3^. However, the nature, timing and contributions of other potential bacterial partners^4–6^ or the role of interactions with viruses^7–9^ remain contentious. To address these questions, we employed advanced phylogenomic approaches and comprehensive datasets spanning the known diversity of cellular life and viruses. Our analysis provided an updated reconstruction of the last eukaryotic common ancestor (LECA) proteome, in which we traced the phylogenetic origin of each protein family. We found compelling evidence for multiple waves of horizontal gene transfer from diverse bacterial donors, with some likely preceding the mitochondrial endosymbiosis. We inferred plausible traits of the major donors and their functional contributions to LECA. Our findings underscore the contribution of horizontal gene transfers in shaping the proteomes of pre-LECA ancestors and hint to a facilitating role of Nucleocytoviricota viruses. Altogether, our results suggest that ancient eukaryotes originated within complex microbial ecosystems through a succession of diverse associations that left a footprint of horizontally transferred genes.

## Main text

The sharp divide in terms of cellular complexity between eukaryotes and prokaryotes has been deemed “the greatest single evolutionary discontinuity to be found in the present-day world”^10^, and the possibility of extensive gene transfer through prokaryotic (endo)symbiosis has long been considered a stepping stone in this process^11^. The current consensus on eukaryogenesis revolves around scenarios that always involve an endosymbiotic relationship with extensive gene transfer between an alphaproteobacterial endosymbiont and a host with an Asgard archaeal ancestry^2,3^. Phylogenomics has shed light on the phylogenetic placement and potential traits of these two established partners^12–16^. However, some proposed eukaryogenesis models involve at least one additional partner^5^, or even serial interactions with various non-alphaproteobacterial symbionts acting as gene donors^4,6^. The large dominance of bacterial over archaeal contributions in reconstructed LECA proteomes^4,17^, and the observation that only a small fraction of bacterial-derived proteins can be confidently traced back to Alphaproteobacteria, may hint to additional bacterial contributions^4,18^. However, alternative explanations to such observations include the difficulty of reconstructing ancient phylogenetic relationships and the presence of horizontal gene transfer (HGT) carried by the Asgard archaeal or alphaproteobacterial partners.

Here we asked the objective question of whether protein families within LECA could trace back to ancestors different from Alphaproteobacteria or Asgard archaea with a level of support similar to those tracing to these broadly-accepted partners. To alleviate potential artefacts arising from phylogenetic reconstruction, including the effects of unsampled lineages, contamination, and recent HGT, we used state-of-the-art methodologies and compiled curated datasets including representative proteomes with the highest possible qualities, from which we purged low-quality sequences, recent paralogues and sparsely distributed proteins. Our results uncover at least two major signals of bacterial ancestry different from Alphaproteobacteria, and a consistent set of gene acquisitions inferred to be mediated by Nucleocytoviricota viruses.

### Reconstruction of the ancestral eukaryotic proteome

To leverage the recent explosion of genome data across the tree of life, particularly among eukaryotes, we re-inferred the LECA proteome using an automated approach similar to previous studies^4,19–21^. To minimise known methodological issues in homology and phylogenetic inference, we subsampled existing data to obtain a balanced representation across the eukaryotic tree of life (eTOL), while ensuring a tractable size and the highest possible quality (**see Methods, Supplementary Methods, Supplementary Tables 1 and 2)**. We additionally curated the selected proteomes to remove low-quality and low-complexity proteins. Given our focus on deep evolutionary nodes, we kept a single representative of clusters of recent eukaryotic in-paralogs. We replicated this procedure to generate three alternative 100-proteome datasets (eTOLDB-A, -B, and -C) that overlap in about 46% of their proteins (**Supplementary Fig. 1)** and allow assessing the data-dependency of our results. We clustered proteins in these datasets into orthologous groups (OG) and defined putative descendants of LECA (LECA-OG) those OGs containing at least five different species, and at least three out of nine eukaryotic supergroups and the two main eukaryotic stems after removing potential contaminants (**see Methods and Supplementary Fig. 1 and 2**). LECA-OGs were highly consistent across datasets (>96%). To further refine these families we used protein alignment profile similarity searches against a broad database (broadDB) comprising order-level prokaryotic pangenomes reconstructed from over 65,000 genomes available at GTDB^22^ and sequence representatives of over 1.3 million clusters of viral sequences^23^ (**see Methods**). This approach ensured maximal coverage of extant diversity while minimising the impact of database biases and recent HGT. We next reconstructed Maximum Likelihood (ML) phylogenies from the LECA-OG expanded with the closest broadDB hits (**see Methods**). This served to assess the monophyly of the eukaryotic proteins, which were split into different, monophyletic LECA-OGs (mLECA-OGs) if necessary. We repeated the same procedure with this new set of mLECA-OGs by building new alignment profiles, and repeating the broadDB search and phylogenetic reconstructions (**see Methods**). This resulted in a final set of mLECA-OGs (with 79% consistency across datasets) and their phylogenies in the context of their closest non-eukaryotic homologs. Analysis of the earliest splits in the mLECA-OG phylogenies indicated that only 3% of the OGs could possibly result from HGT between eukaryotic supergroups (**see Methods**).

Our reconstructed LECA proteomes comprised an average of 12,907 or 7,751 mLECA-OGs (OGs hereafter) depending on the relaxed (3 supergroups) or strict (5 supergroups) criterion (**Supplementary Table 3**), which is in line to previous recent estimates (e.g. 10,233 in^21^). We approximated the functional potential of LECA by annotating these OGs using a taxonomically-delimited set of the KEGG Orthology database (KOs) (**see Supplementary Methods**) and then we inferred a relaxed consensus annotated proteome of 5,317 KOs (consensus proteome hereafter, see the Zenodo repository for the list of KOs) consistently predicted in two or more datasets, or in one dataset and the strict criterion (**see Methods**). Given the limitations of an automated approach (**see Supplementary Discussion),** the purpose of this reconstruction is to approximate gene families and functional categories likely present in the ancestral LECA proteome, rather than to provide a detailed picture of LECA’s traits.

To assess the completeness and validity of our reconstructed LECA proteome, we assessed the reconstructed functional potential in terms of its congruence with previous inferences about LECA. Our reconstruction confirms earlier studies depicting a complex LECA, covering the main features of extant eukaryotes^24^ (**Fig. 1a and b, and Supplementary Discussion**). LECA had the core metabolism and machinery for nucleic acid and protein processing, endocytosis, and the processing of extracellular particles, and contained phagosomes, lysosomes and peroxisomes, as well as mitochondria with the ability of aerobic respiration, heme and Fe-S cluster assembly, among other pathways. We infer a cytoplasm similar to that of extant eukaryotes with a cargo-based transport using the motor proteins (dynein, dynactin and kinesin). Conversely, the cell cycle appears rather incipient, with the presence of the basic machinery and a very limited regulation (see **Supplementary Discussion**). Regarding metabolism, the low completeness of the autotrophic Wood–Ljungdahl and the Arnon–Buchanan cycle suggest a heterotrophic lifestyle. Moreover, the reconstruction lacks enzymes related to anaerobiosis (Pyruvate:ferredoxin oxidoreductase –PFO–, [FeFe] hydrogenase and enzymes involved in the rhodoquinone synthesis), suggesting that LECA was most likely an aerobe (**Supplementary Table 4**, **see Supplementary Discussion**). We compared the reconstructed consensus proteome with the proteomes of representative extant free-living unicellular phagotrophs (FLUPs), osmotrophs (FLUOs), and autotrophs (FLUAs) sampled across the eTOL (**Supplementary Table 5**). Although FLUPs share the inferred trophic lifestyle of LECA^2^, we did not observe clear differences in terms of the absolute number of LECA KOs (**Fig. 1c**) and COG categories inherited by osmotrophs and phagotrophs (**Extended Data Fig. 1, Supplementary Table 5**). Amid a wide range of individual variation, specific discrepancies in the relative abundance of functional classes indicate post-LECA adaptations (**Extended Data Fig. 1**). Notably, all trophic groups have significantly expanded repertoires of signal transduction proteins, suggesting post-LECA diversification in ecological and environmental interactions.

**Fig. 1.**
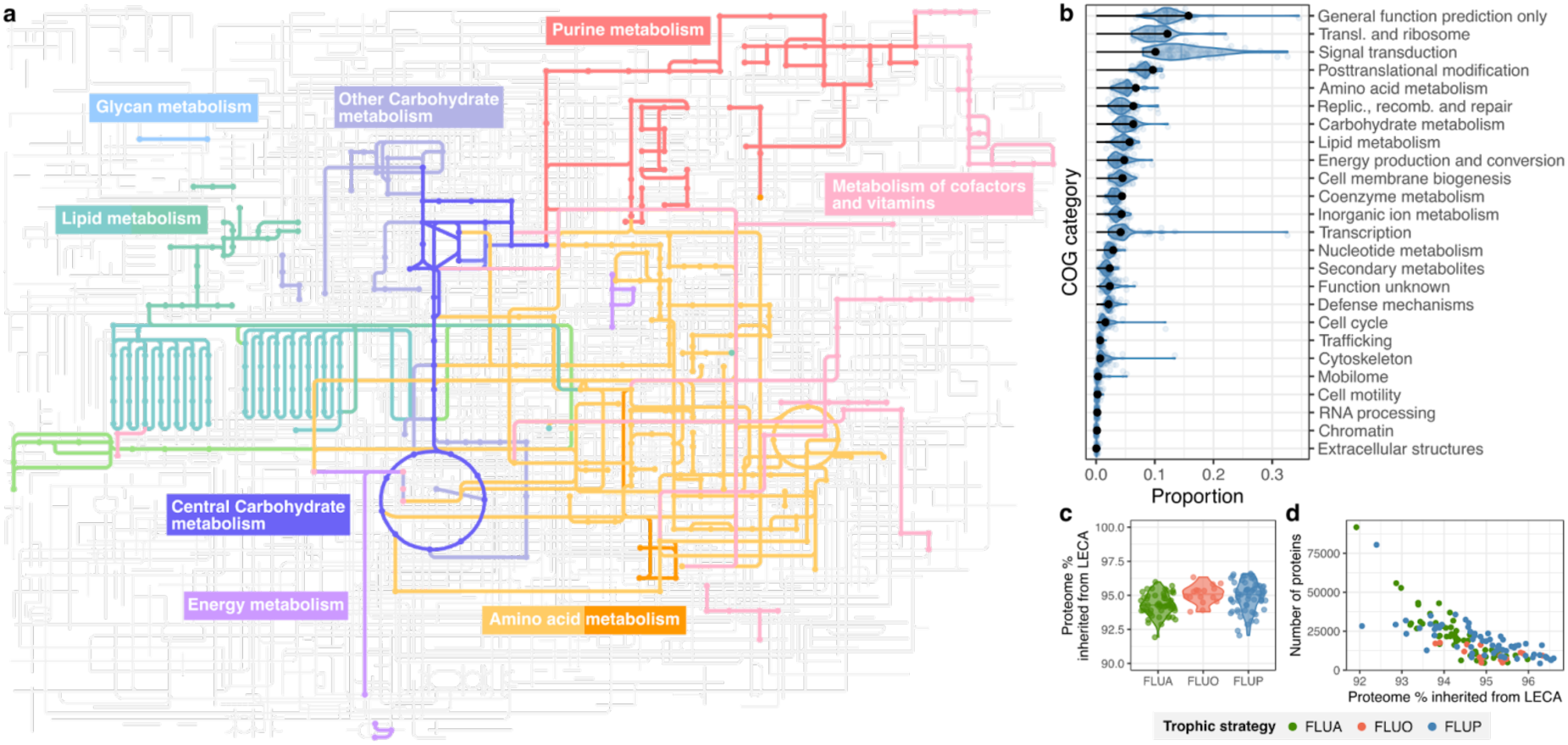
Reconstruction of the ancestral proteome of LECA and its metabolic features. a) Broad overview of the reconstructed LECA metabolism. In light grey: the metabolic map of KEGG. In colors: the modules inferred to have been present in LECA using the consensus proteome, which accounts for the KOs present in at least 2 LECA reconstructions of different eTOLDBs or supported by a LECA group with five eukaryotic supergroups. Each colour refers to a metabolic category according to the coloured boxes. Detailed maps of different modules are available at the Zenodo repository accompanying this article. b) Frequency of COG database functional categories in the consensus LECA proteome (black) and in 63 free-living unicellular phagotrophs (FLUPs, distribution in blue). c) Percentage of KEGG orthologs (KOs) shared between the ancestral proteome of LECA and extant free-living unicellular autotrophs, osmotrophs and phagotrophs (FLUAs, FLUOs and FLUPs, respectively). d) Relationship between the percentage of the proteome inherited from LECA and the extant genome size.

### Distilling the ancestries of the LECA proteome

We next investigated the topologies of the reconstructed trees for each OG and inferred the phylogenetic sister group as the best proxy for ancestry (**see Methods, Supplementary Fig. 3**). A significant fraction of OGs (33%) had only eukaryotic homologs passing our filters, and we classified them as putative innovations (**Fig. 2a)**. Note that such putative innovations may include some protein families with protein domains or KOs annotations present in prokaryotes, as domain reshuffling or extensive sequence divergence may result in failure of homology detection by our procedures. Considering only KOs in the consensus proteome predicted exclusively as innovations, and annotated with eukaryote-exclusive and taxonomically-broad KOs, we compiled a list of 918 KOs that could be considered an updated set of Eukaryotic Signature Proteins (ESPs) sensu ref.^25^ **(Supplementary Table 6)**. For OGs having non-eukaryotic homologs (i.e. acquisitions, ∼53% on average, **Fig. 2a**), we found a diversity of taxonomic assignments, in line with earlier findings ^4,17,18,21,26^.

**Fig. 2.**
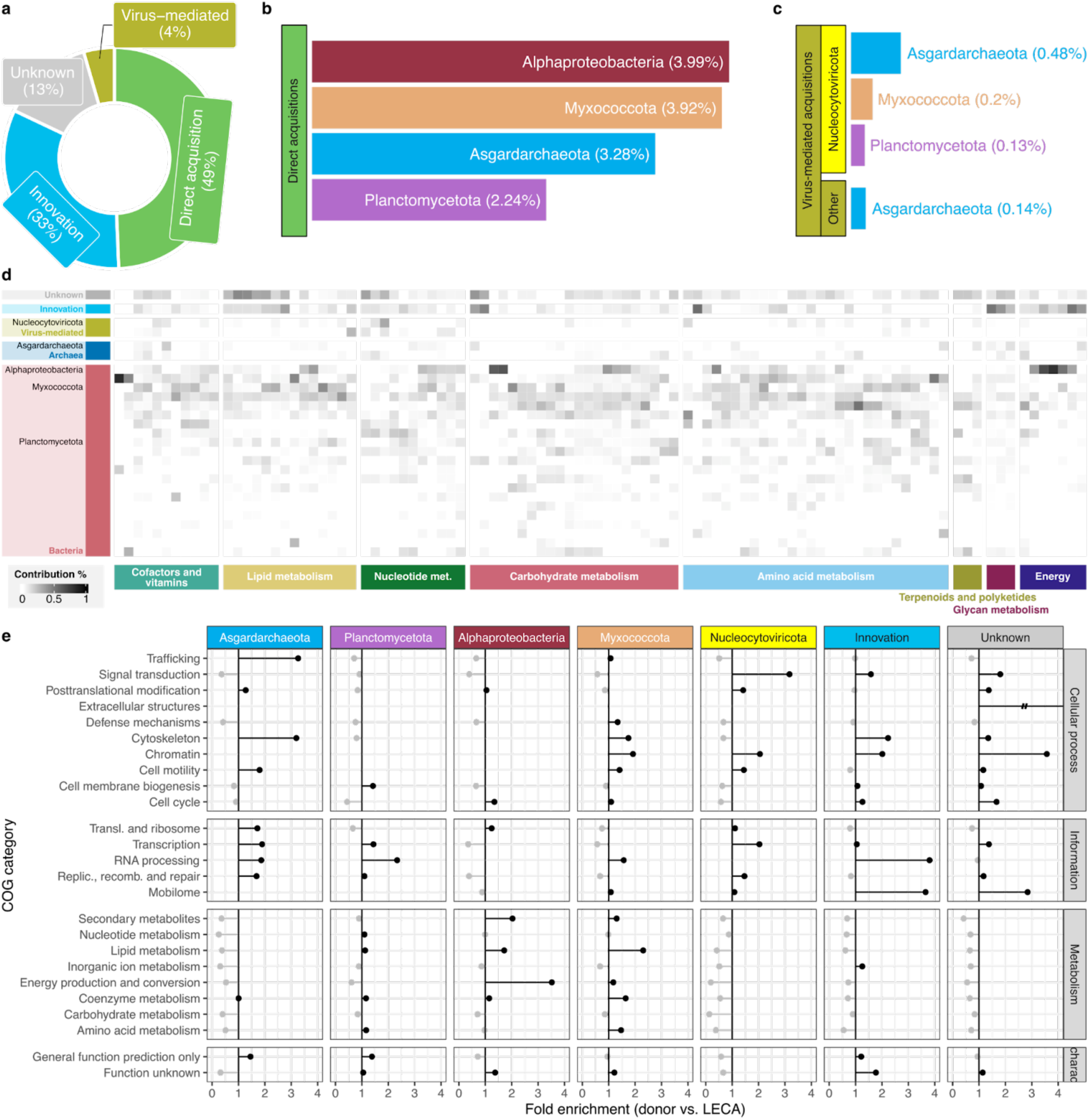
Summary of the LECA ancestries. a) The circular diagram shows the average representation over the LECA proteome across eTOLDBs of innovations, direct- and virus-mediated acquisitions, and families of unknown origin. b) Distribution of the ancestries of the LECA proteome. The barplot shows the main inferred prokaryotic affiliations of genes directly acquired (from prokaryotes to LECA) and their average percentage of representation over the LECA inferred proteomes. c) Shows the same barplot as b but for the virus-mediated acquisitions (from prokaryotes to viruses to LECA). d) Heatmap showing the proportion of the contribution of each inferred ancestry (rows) to LECA metabolic modules (columns). The intensity of the colour shows the percentage of annotated KOs from a given metabolic module that are from the taxa ancestry indicated in the rows (for details, see **Supplementary Table 4** and the data repository). e) Fold enrichment of functional contributions of the non-eukaryotic genes to the LECA consensus proteome. This is obtained by dividing the proportion of transferred genes with that ancestry in that COG category, by the proportion represented by that category in LECA. Therefore, values above 1 mean that the gene is more frequent among the donor’s contributions than in LECA.

As some affiliations may result from phylogenetic noise, and we were interested in identifying potential relevant partners, we assessed the impact of different stringency thresholds on the proportion of trees assigned to the generally accepted main LECA contributors: Alphaproteobacteria and Asgard archaea (**see Supplementary Discussion, Supplementary Fig. 4a)**. Using the thresholds that maximized the proportion of these bona-fide affiliations among selected trees, we identified two consistent bacterial affiliations beyond Alphaproteobacteria (3.99% of the LECA proteome, on average) and Asgard archaea (3.28%) that kept a non-negligible proportion of genes: namely Myxococcota (3.92%) and Planctomycetota (2.24%, **Fig. 2b**). Additional minor bacterial affiliations contributing fewer genes were also identified (**Supplementary Fig. 5**), including Chloroflexota and Gammaproteobacteria consistent with previous observations^4^, however these failed to pass the thresholds across all datasets and were therefore not considered in subsequent analysis. The overall representation of the chosen major taxonomic assignments remained highly similar across datasets and stringency criteria, indicating this result is independent of the eTOLDBs composition (**Supplementary Fig. 3**). Additionally, we found that trees pointing to these alternative taxonomic assignments were not significantly more difficult to resolve than those with alphaproteobacterial sister groups **(Supplementary Fig. 6)**. From these analyses, we conclude that phylogenetic noise alone cannot result in the observed diversity of taxonomic assignment patterns.

Given the presence of putative additional bacterial donors, we considered the possibility that they could result from wrong phylogenetic placement due to incomplete sampling of Alphaproteobacteria. That is, in the absence of a close alphaproteobacterial relative of a given gene in our database, this gene might be assigned to a different bacterial group. We assessed this possibility by applying our OG reconstruction and phylogenetic assignment procedure to Alphaproteobacterial genomes, while removing this group from the BroaDB (**see Methods**). Expectedly, this “negative control” shows a majority of trees assigned to the closest sister of the missing taxon, Gammaproteobacteria, followed by alternative taxonomic assignments with abundances that decrease as they correspond to more distant relatives (control bars in **Supplementary Fig. 4b**). This expected pattern is not observed for our Myxococcota or Planctomycetota affiliations, supporting they represent acquisitions from donors distinct from the mitochondrial ancestor (see different trends in control and LECA bars in **Supplementary Fig. 4c** and **Supplementary Discussion**).

Notably, approximately 4.5% of the mLECA-OG phylogenies show a viral sister group (**Fig. 2a**), with Nucleocytoviricota, a phylum containing giant viruses, being the most frequent viral sister (74%), resulting in 3.32% of the LECA proteome. Trees containing viral sister groups were further analysed in search of the first non-viral sister, which detected putative virus-mediated acquisitions from diverse prokaryotes (**Fig. 2c**, see **Methods** and **Supplementary Discussion**). In addition to innovations and acquisitions (either direct or virus-mediated), a constant fraction of the trees (∼13%, on average) had an uncertain taxonomic origin or acquisition directionality, due to a poor representation of the non-eukaryotic sequences in the tree, and we labelled these as of “unknown origin”. Consistently, signals for all the major prokaryotic affiliations identified here have also been detected to different degrees by previous large-scale studies^4,17,18,21,26^, which used far less complete eukaryotic and prokaryotic genome databases and did not include viruses. Database completeness rather than methodological choices are likely to drive differences across studies, as supported by a re-analysis of the phylogenies of one of these studies with the analytical procedures used here (see **Supplementary Discussion**, and **Supplementary Fig. 7**).

We next assessed the predicted functions of acquisitions assigned to different taxonomic affiliations, and found generally broad and patchy distributions across the reconstructed LECA metabolism (**Fig. 2d**). Conversely, most metabolic modules and cellular processes showed contributions from innovations and from acquisitions of different affiliations, suggesting a high degree of chimerism in LECA (see **Supplementary Discussion, Supplementary Fig. 8, Supplementary Table 7**). Nevertheless, some functional categories show over-representation of certain origins (**Fig. 2e**), confirming earlier described trends such as the information/metabolism divide in terms of archaeal and bacterial ancestries^26^ or alphaproteobacterial origins of energy production (see **Supplementary Discussion**).

### Relative timing and potential metabolic nature of gene donors

To investigate whether genes showing different ancestries might have been acquired in the form of “transfer waves” (i.e. when the inferred timings of different HGTs cluster around a similar period), we applied a previously described branch length ratio approach^4^, as recently implemented into a Bayesian inference framework^27^. This approach measures lengths of the branch separating the ancestor of the LECA group from the most recent common ancestor of its non-eukaryotic sister group, normalised with the median LECA-to-tip branch length within the LECA clade, which refers to the same geological time (from LECA to present) and therefore provides a relative time for the acquisition (see **Methods**). In this framework, longer normalised lengths are interpreted as more ancient relatedness.

Our results for the donor lineages (**Fig. 3a, Extended Data Fig. 2**) depict distinct branch length distributions peaking at different values, suggesting different relative timings for the acquisitions. With slight variations, we detected consistent trends in the relative ordering of the distributions across datasets and stringency levels (**see Extended data Fig. 3**). First, Asgard archaea is the most ancestral signal. Among the bacterial donors, Planctomycetes tend to show the longest stem lengths, suggesting an early interaction, followed by later contributions from the mitochondrial ancestor and Myxococcota, with largely overlapping transfer waves. Notably, all identified donor bacterial taxa are common components of microbial mats, and the inferred order of acquisition of Asgard archaea, Planctomycetota and Alphaproteobacteria is similar to the relative depth in which each of these taxa is found in a deeply-characterized mat^28^ (**Fig. 3b**).

**Fig. 3.**
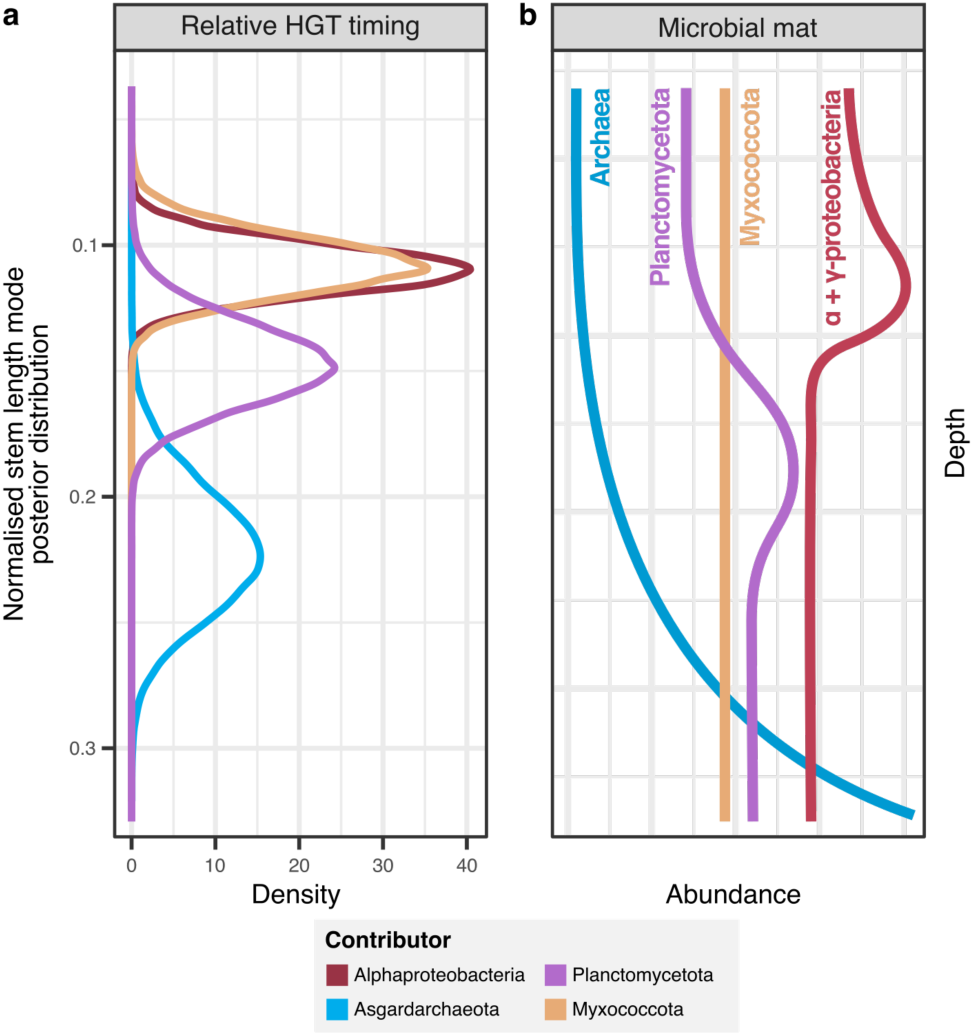
LECA waves of acquisition in the context of microbial mats. a) Posterior distribution of the mode (distribution peak) for the normalised stem lengths of each putative donor. Shorter stem lengths would refer to more recent transfers while longer would indicate earlier acquisitions. The colours indicate the different donors to LECA according to the legend at the bottom. The posterior distributions for each database and criterion are available in the **Extended Data Fig. 2**, which also includes the non-donor signals Gammaproteobacterial and Thermoproteota. b) Schematic relative abundances of the inferred donors across the different layers of a microbial mat according to ref. ^28^.

We used the above branch length distributions to refine the set of proteins likely acquired from the same donor in a single wave by removing protein families with anomalous branch lengths (see **Methods**). We then inferred the metabolic capacities possibly present in each considered donor by assessing metabolic pathways commonly present and complete among the extant genomes from each donor clade or category that also shared more than 50% of the LECA’s KOs assigned to this ancestry (donor’s descendant genomes, **see Methods**). For comparison, we also considered more restrictive reconstructions by constraining the search to genomes of the currently assumed ancestor of the archaeal lineage in eukaryotes (Heimdallarchaeia)^12,29^. These reconstructions, which expand KO acquisitions in LECA with commonly co-occurring KOs in extant organisms of that donor clade, provide a first indirect approximation for the core set of metabolic pathways possibly present in the ancestral donors (**Fig. 4**, **Extended Data Fig. 4 and 5, Supplementary Data**). Note that this reconstruction is different from standard ancestral state reconstruction, which necessitates higher resolution on the donor’s taxonomic affiliation and the underlying species phylogeny. Instead, our approach relies on the assumption that KO co-occurrence patterns commonly observed in extant genomes of one phylum were likely in place in an unknown ancestor of that phylum, which transferred certain functions to LECA. The discussed inferred pathways maintained their overall tendency when we altered the database composition with an iterative resampling approach, penalizing over-represented taxa **(Supplementary Methods)**.

**Fig. 4.**
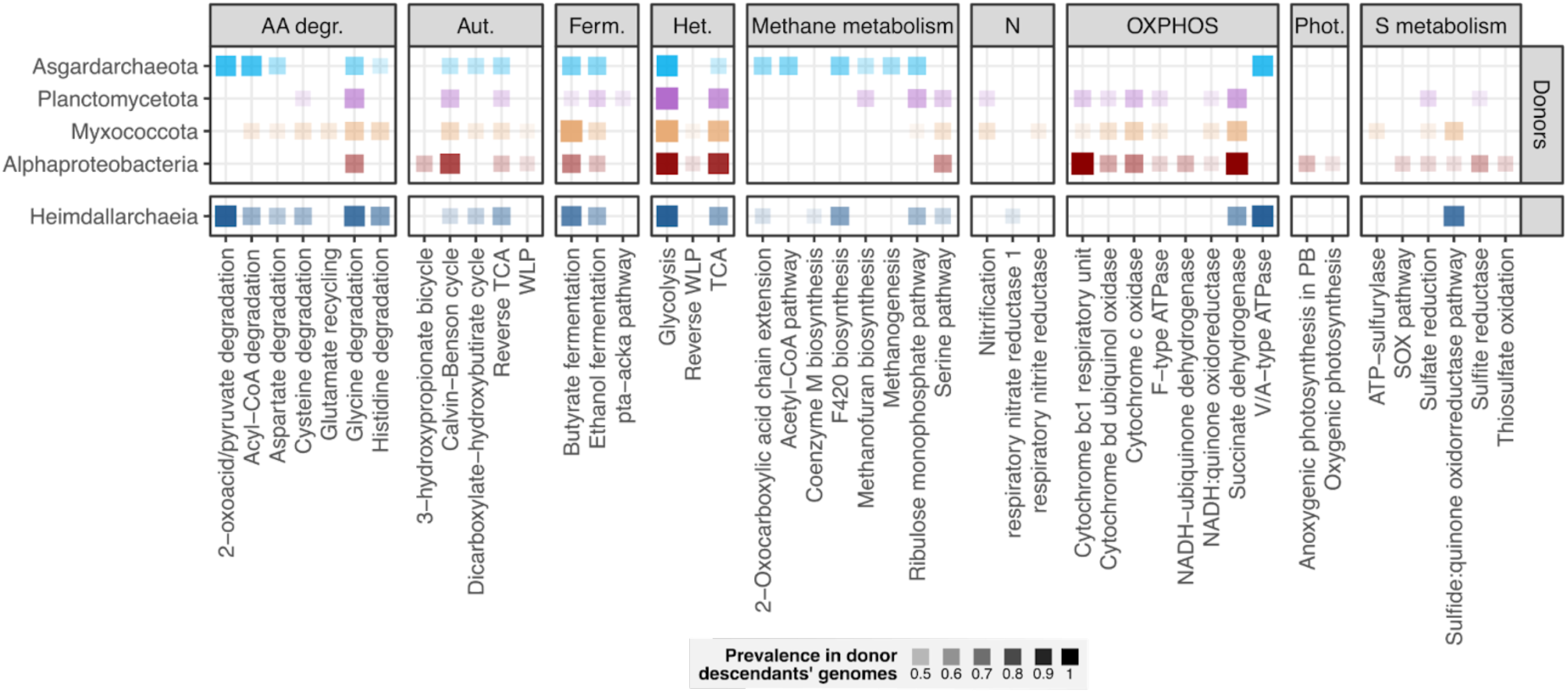
Potential metabolic features of the prokaryotic donors. Prevalence of some metabolic features for each donor. The opacity of the square (with colours congruent with Fig. 2) indicates the proportion of genomes with at least 50% of the metabolic pathway completeness. Note that we include the currently proposed Asgard sister to eukaryotes for comparison (Heimdallarchaeota, at the bottom). Pathways and KOs with more than 0.4 prevalence are shown. A detailed version of this plot, with the prevalence for all the metabolic pathways and KOs assessed, is available in the **Extended Data Fig. 4**. The equivalent plot considering the origin in LECA of genes, from each donor, involved in these metabolic features is available in the **Extended Data Fig. 5**. Abbreviations: “AA degr.” amino acid degradation, “Aut.” autotrophy, “Ferm.” fermentation, “Het.” heterotrophy, “N” nitrogen metabolism, “OXPHOS” oxidative phosphorylation, “Phot.” photosynthesis, “S metabolism” sulphur metabolism.

Considered altogether, these observations depict a complex scenario involving a series of HGT waves from a diverse set of donors into the proto-eukaryotic lineage, as proposed by the pre-mitochondrial symbioses hypothesis^6^. The inferred taxonomic identity, metabolic features, and order of acquisition also support previous ideas of syntrophic origins within microbial-rich environments, such as microbial mats^30^ (**Fig. 3b**), but additionally provide remarkable new insights. For instance, while the Hydrogen Sulphur-transfer-based (HS) syntrophy hypothesis positioned Deltaproteobacteria (which includes Myxococcota) as the eukaryogenesis host^5^, our analysis suggests Planctomycetota is an earlier contributor. This, together with the notable cellular complexity of this group^31^, including the remarkable potential for cell engulfment observed in one genus^32,33^, may lend some support to consider Planctomycetota as an alternative bacterial host, as previously debated^34,35,36^. Nevertheless, we note that our data does not provide information on the host or endosymbiotic nature of any partner.

As to the potential metabolic interactions, the incomplete and indirect nature of our reconstructions makes it difficult to favour a specific proposal, but provide information to contrast different proposed models. For instance, our results discourage a methanogenic nature of the Asgard ancestor as proposed in various of the earliest models^5,30,37^, as none of the Asgard archaeal genomes that resemble the LECA footprint have a complete methyl-coenzyme M reductase (MCR) and coenzyme M methyltransferase (MtbA) **(Supplementary Discussion).** There is support for a fermentative metabolism (with 0.63 prevalence for ethanol fermentation and 0.67 butyrate fermentation in the archaeal ancestor, see **Extended Data Fig. 4**) as suggested in the reverse-flow model^38^. Our inference moderately supports a potentially photosynthetic mitochondrial ancestor (with prevalence for the different photosynthetic systems ranging from 0.34 to 0.51), as proposed earlier^5,39,40^. Sulphate reduction is inferred with certain support for the Myxococcota donor as proposed in the revised HS syntrophy hypothesis^5^ (0.44 prevalence for sulphate reduction) but also in the Planctomycetota donor (0.5 prevalence), suggesting the model could be adjusted to this alternative host to accommodate our relative timing observations. Both cultured Asgard archaea grow in syntrophy with sulfate-reducing Desulfobacterota partners^41,42^, and sulfur-based interactions have been hypothesized between extant Asgard archaea and Planctomycetota in mangrove sediments^43^, suggesting this type of syntrophy can be established between Asgard archaea and diverse bacterial groups. We also observe a weak potential for ammonium oxidising (anammox) metabolism in the Planctomycetota donor (prevalence of 0.23). Importantly, the reconstruction of the potential metabolic features present in the mitochondrial ancestor strongly supports the presence of aerobic respiration but lend no support to the presence of Fe-Fe-Ni-Fe- and Fe-only hydrogenases as implied by the Hydrogen hypothesis or the reverse-flow models^30,37,38^. Nevertheless, further phylogenetic, genomic, ecological and biochemical research is needed to elucidate the nature of these early interactions.

### Viruses as mediators of gene acquisitions during eukaryogenesis

The contribution of viruses in shaping the proteome of LECA has been discussed in earlier studies^44–46^. A recent analysis surveying viral sequences across eukaryotes identified a significant amount of virus-to-eukaryote transfers across the eTOL, despite a dominance of eukaryotic-to-virus transfers^8^. Our analysis provides a more systematic and specific assessment of potential virus-mediated HGT into LECA, which we found to account for at least 4.5% of all acquisitions, after strict filtering to ensure timing and directionality. Importantly, most such transfers (74%) were found to be mediated by a specific type of viruses: Nucleocytoviricota. Nucleocytoviricota, also referred to as nucleocytoplasmic large DNA viruses (NCLDVs), is a phylum of viruses known to infect diverse eukaryotes and to include giant viruses with large genomes^47^. We explored trees that showed Nucleocytoviricota as the first sister to LECA and included farther prokaryotic sister groups, as their topology suggested indirect acquisition of prokaryotic genes by LECA through viral mediation. These prokaryotic sisters contained, among other prokaryotic groups, all of the donor groups detected in the previous analysis, with the most represented donor corresponding to Asgard archaea (∼10%, **Fig 2c, and Supplementary Table 8**). Functional analysis of viral-transferred genes indicated over-abundance of genes related to signal transduction mechanisms, mainly kinases, chromatin, and post-translational modification proteins (**Fig. 2e**). This observation is compatible with other findings indicating this type of viruses infected proto-eukaryotes^44^, and opens the possibility of viral-mediated transfers to the LECA lineage from co-existing but now-extinct, eukaryote-like lineages, including genes of prokaryotic origin previously acquired by these lineages^48^.

## Discussion

Our results confirm and expand earlier results supporting significant gene flow from diverse prokaryotic ancestors preceding LECA^4,21^, and uncover a role for viruses as potential mediators of such transfers. Considering the distributions of phylogenetic distances to the closest relatives, our data support pre-mitochondrial waves of gene acquisitions from additional major prokaryotic donors as the most likely scenario^6,49^, against a backdrop of continuous HGT contributions from a diversity of other minor sources. We find support for at least two major non-proteobacterial bacterial donors transferring a significant amount of genes: Planctomycetota, and Myxococcota. Transfers from these donors have been identified in earlier studies^4,18,21^, including small-scale detailed ones such as the origin of some steroid biosynthesis enzymes from Myxococcota^50,51^. Our results uniquely support the existence of sizable genetic exchange with these alternative donors, hinting at stable interactions beyond those between the Asgard archaeal ancestor and the alphaproteobacterial ancestor of mitochondria. Although different forms of interactions beyond symbiosis could explain this genetic exchange our results support that current eukaryogenesis models involving a single interaction between an Asgard archaeal ancestor and the alphaproteobacterial ancestor of mitochondria should be expanded to cover additional interactions with other relevant partners. Models involving interactions with additional bacterial partners could be refined in light of the potential donors identified here^5,6^. Our results also uncover a role of viruses, particularly Nucleocytoviricota, as mediators of some of these prokaryotic gene transfers. This finding can be considered in the light of previously proposed roles of viruses during eukaryogenesis (see **Supplementary discussion**).

Our relative timing estimates suggest early interactions between an Asgard archaea and a Planctomycetota, followed by later contributions of Myxococcota and the acquisition of the mitochondrion. Thus, the prokaryotic-to-eukaryotic transition was likely a gradual and complex process, with a constant flow of HGT from diverse sources. Although our data does not inform on the potential type of the interactions (endosymbiotic, ectosymbiotic, parasitic, etc), the inferred metabolic capabilities of the detected donors can be used to contrast existing metabolic scenarios for eukaryogenesis, adjust them, or propose novel ones. Microbes are known to form complex communities such as microbial mats or complex biofilms^52^, where viruses are also an active part, and it is reasonable to consider that the ancestors of LECA lived in such complex environments. Successive, perhaps overlapping, long-term interactions with different microbes and the mediation of viruses would have led to extensive gene transfer, progressively shaping the chimeric proteome of LECA.

Altogether, our results challenge models depicting eukaryogenesis as a one-off event resulting from a binary symbiotic interaction, and support a paradigm shift towards more gradual scenarios in which the prokaryotic-to-eukaryotic transition was achieved through serial interactions with a diverse range of bacterial partners leading to significant gene transfer, and with viruses playing a role as mediators.

## Methods

### 1. Data retrieval, curation and proteome database reconstruction

#### 1.1. Eukaryotic Tree of Life database (eTOLDB) construction

To guide dataset selection and LECA reconstruction, we used the phylogenetic backbone of the eukaryotic tree of life (eTOL) and the nine eukaryotic supergroups classification depicted in (**Supplementary Fig. 1a**). This is a consensus between recent eTOL reconstructions^53–57^ and an operational definition of two stems and nine supergroups that largely reflect previous classifications^58^. Additional details are provided in (**Supplementary Methods**).

We manually selected and retrieved 276 eukaryotic proteomes (**Supplementary Table 1 and Supplementary Data**) downloaded from NCBI^59^, EukProt^60^, P10K^61^,UniProt^62^, Ensembl^63^ and SGD^64^, aiming to cover all supergroups represented in our reference phylogenetic backbone (**Supplementary Fig. 1**). We aimed at a balanced representation of the divisions and prioritised the most complete and highest quality genomes for each considered division based on information available in the hosting databases. For each proteome, we only kept the longest isoform per gene, as indicated in the GFF files or FASTA headers. The isoform-free proteomes were then submitted to an analysis of completeness using BUSCO v5.5^65^ with default parameters and the eukaryotic_odb10, and we extracted basic statistics using seqkit stats v2.8.2^66^. To assess for the presence of uncertain amino acids (Xs) and low complexity regions in the proteomes, we ran SEG^67^ with default parameters. We used an in-house script (see **Code availability**) to trim the low-complexity ends of the proteins and calculated the coverage of low-complexity regions in the protein. We considered a low-complexity protein if the coverage of low-complexity regions was higher than 25% of the protein and discarded it for the downstream analyses. Moreover, we calculated the proportion of low-complexity proteins in the whole proteome as a measure of the quality of the protein set. We also discarded proteins that exceeded the 10,000 amino acids or were smaller than 50 amino acids. This resulted in a set of clean proteomes. To remove redundancy and recent duplications (with respect to eukaryotic evolution), we clustered each proteome individually using MMseqs v2^68^ with strict thresholds: minimum amino acid identity percentage of 80%, minimum target coverage of 0.5 and coverage mode 2 (--min-seq-id 0.8 -c 0.5 --cov-mode 2). The representative sequences of the clusters constituted the final proteome. To assess the effect of the cleaning and clustering steps, we ran again BUSCO v5.5 and seqkit stats v2.8.2. We discarded the proteomes that suffered a combined reduction in the number of proteins higher than 50%, having a low-complexity proteins percentage higher than 50% and a BUSCO completeness reduction higher than 30%. This resulted in 30 proteomes being excluded, and we continued with the 256 remaining proteomes.

We built three eTOL datasets of 100 proteomes each, which we named eTOLDBA, eTOLDB and eTOLDBC. For this, we first classified the 256 proteomes based on taxonomy and distributed them in nine different supergroups and 28 divisions (**Supplementary Fig. 1**). When less than four species were available for a division, we included all in the three eTOLDB databases. For the remaining divisions we selected different, partially overlapping, species sets for each eTOLDB. eTOLDBA contained, whenever possible, the most complete proteomes and the fewest parasites. More incomplete proteomes were added in turn in eTOLDBB and in eTOLDBC. Proteomes for model species *Homo sapiens, Arabidopsis thaliana* and *Saccharomyces cerevisiae* were included in the three databases despite being part of large taxonomic groups. This was done for later annotation purposes. The list of chosen species for each dataset can be found in **Supplementary Table 1**. Finally, a total of 185 species were present in the three eTOLDBs with an overlap of 58 species between eTOLDBA and eTOLDBB, and 53 species between eTOLDBA - eTOLDBC and eTOLDBB - eTOLDBC. This resulted in an average protein overlap of 46% (see **Supplementary Fig. 1)**.

#### 1.2. Broad non-eukaryotic database construction (BroadDB)

To ensure an even sampling of the prokaryotic genome repertoire, while minimising the impact of intra-domain HGT as a confounding factor of prokaryotic ancestry, we downsampled available prokaryotic genomes using a pangenome approach. For this, we took all species representatives in GTDB r207^22^ (62,291 bacteria and 3,412 archaea, see and **Supplementary Data**) and selected one genome per genus (*sensu* GTDB), choosing the one with the maximal CheckM^69^ completeness and, in case of ties, minimal CheckM contamination. The median CheckM completeness was 94.50% and the median contamination was 0.990%. The proteomes of each genus representative were then grouped at order level and clustered with mmseqs2 easy-linclust^68^ (--min-seq-id 0.3 -c 0.5 --cov-mode 0 --cluster-mode 2 -e 0.001). We kept a sequence representative for clusters present in 15% or more of genus representatives in that order, as 15% is the cut-off criteria for the “cloud” component of the pangenome^70^. To this dataset we added representative sequences from 1,319,927 viral protein clusters at 95% identity percentage available at RVDB v25.0^23^ (see **Supplementary Data**). For taxonomic assignment of the sequences, we follow the GTDB and RVDB taxonomies throughout the manuscript.

### 2. Reconstruction of LECA

#### 2.1. LECA-Orthogroup (LECA-OG) reconstruction

We ran OrthoFinder v2.5.5^71^, using BLAST v2.15.0+^72^ for the homology search with the filter for low complexity (-seg yes). Before the clustering step, BLAST results were filtered to remove hits with an e-value above 1e-05 and an overlap below 0.5. We additionally removed hits with disparate sizes (when hit was >2.5 longer or <0.5 shorter than the query) to avoid artefactual clustering of families due to domain sharing. The clustering part of OrthoFinder was executed using two different inflation parameters (I = 1.5 and I = 3.0). Comparison between the two datasets showed that I = 3.0 gave 30,000 more groups on average than I = 1.5. When filtering to retain only LECA groups then there was an increase of 990 groups on average between I = 3.0 and I = 1.5. Given the small difference we chose to continue with the larger groups (I = 1.5) as they could be split based on tree topology at a later point.

Orthogroups (OGs) as defined by OrthoFinder were filtered to select those likely present in LECA (LECA-OG) based on their distribution across the eTOL. LECA-OGs had to comprise proteins from at least five species, and have representatives of the two stems and three or more different supergroups for a relaxed definition of LECA and five or more supergroups for a strict definition of LECA. These same criteria for the identification of LECA were used in downstream analyses.

#### 2.2 Contamination assessment and filtering of the LECA-OG sequences

To assess potential non-eukaryotic contaminant proteins, we followed a taxonomy-based prok-euk contamination detection. First, we built a broad database to assess contamination. This database contains all prokaryotes from the reference release of GTDB r222^22^, all eukaryotic proteomes in UniProt^62^, EukProt^73^ and a selection of the P10K^61^ project proteomes. To reduce redundancy and size of the database, we clustered the proteomes of each class using mmseqs v2^74^ with the following thresholds: 75% identity and 66% coverage. To flag the proteins as potential contaminants, we followed the approach described in Williamson et al.^57^. We performed a diamond blastp v2.1.9^75^ search against a broad clustered database using an e-value threshold of 1e-3, retrieving a maximum of 15 sequences. We considered a protein as contaminant, if 85% of its first 15 hits were prokaryotic, and they showed an identity higher than 85%.

To assess the possibility of eukaryotic contamination, we performed two decontamination strategies: a taxonomy-based, and an identity-based euk-euk contamination detection. For the taxonomy based strategy, we built a database using the eTOLDB sequences and performed an all-vs-all diamond blastp v2.1.9 search. We retrieved the 15 first hits with an identity higher than 85%, and considered contaminants the sequences with 85% of the first hits from the opposite stem (following the taxonomic framework in the Supplementary Fig. 1). We also used an identity-based strategy where sequences from different stems with a blast based identity above 95% were considered contaminants, as we could not establish the directionality of the contamination.

To finally clean putative contaminants within the same LECA-OG, we performed a multiple sequence alignment of the OG and then calculated pairwise identities between all the sequences of the orthogroup belonging to different stems. Those sequences with at least a global 85% pairwise identity were considered contaminants. We removed all the sequences considered contaminants from the LECA-OGs and reconsidered which LECA-OGs passed the LECA criteria with the remaining sequences.

#### 2.3. Expanding LECA-OGs with closest non-eukaryotic homologs

We aligned the sequences with MAFFT v7.525^76^ and built a profile with HMMbuild v3.3.2^77^. We ran hmmsearch v3.3.2 with this profile against the BroadDB described above. Results were filtered using an e-value threshold of 0.1 for both the complete sequence and the first domain found. Then, to ensure that the coverage requirements were maintained we performed a BLASTP v2.15.0+^72^ search for each resulting non-eukaryotic sequence to the database of Eukaryotic sequences from the orthogroup. BLAST results were filtered with an e-value threshold of 1e-05, a coverage threshold of 0.5 over the full length of the non-eukaryotic sequence and ensuring that the target sequence was not 2.5 times longer or 2.5 times shorter than the query sequence. The top-hits that passed these filters were considered the closest non-eukaryotic homologs and were added to the sequences of the corresponding LECA-OG to compute phylogenies (see section below). The number of non-eukaryotic sequences in the trees was limited in the following way: if there were less than 400 eukaryotic sequences, the non-eukaryotic sequences were added until reaching the 500 sequences. When more than 400 eukaryotic sequences were present, up to 100 non-eukaryotic sequences were added. Eukaryotic groups with more than 500 sequences were processed differently. First, genes were assigned a KO (see below) and then sequences from the same OG with the same KO were grouped together and processed as separate OGs.

#### 2.4. Phylogenetic tree reconstructions

We used the PhylomeDB tree reconstruction pipeline^78^ with slight modifications. In brief, we computed a consensus alignment using MergeAlign v3^79^ from the alignments obtained by three different alignment programs (MUSCLE v5.2^80^, MAFFT v7.525^76^ and Kalign v3.4.0^81^) in forward and reverse orientations. This alignment was then trimmed using trimAl v1.4.rev15^82^ with the -gappyout option. Finally, a maximum likelihood (ML) tree was reconstructed using IQ-TREE v3.0.1^83^ and parameters --mset DCmut, JTTDCMut, VT, WAG, LG --madd LG4X, LG4M, LG+C10+R4, LG+C20+R4, LG+C30+R4, LG+C40+R4 –mrate E, G, R, I, G+I, R+I --mfreq FU, F -B 1000 --bnni --alrt 1000 --boot-trees --wbtl. Among the available substitution models, we selected the single matrix models DCmut^84^, JTTDCMut^84^, VT^85^, WAG^86^ and LG^87^ and the mixture models, LG4X, LG4M^88^, LG+C10+R4, LG+C20+R4, LG+C30+R4, LG+C40+R4^89^. We included different rate heterogeneity parameters, the lack of heterogeneity (E), Gamma distributed rates (G), the FreeRate model (R), and their combinations with a proportion of invariant sites (I). We further added two different equilibrium frequency parameters, the one provided by the model (FU) and empirical frequencies (F). ModelFinder was used to select among the combinations of all the models and parameters. Branch support was assessed with 1,000 Ultra Fast Bootstrap replicates optimised by using the nearest neighbour interchange correction (the --bnni option)^90^ and the Shimodaira–Hasegawa approximate Likelihood Ratio Test (SH-aLRT) ^91^.

#### 2.5. Monophyletic LECA orthogroups (mLECA-OG)

For each LECA-OG, we reconstructed a phylogeny following the phylogenetic reconstruction pipeline detailed above. We annotated the tree with the taxonomy of each group and retrieved the largest monophyletic group containing only eukaryotic sequences and fulfilling the relaxed LECA criterion (5 species, 3 supergroups, and 2 stems, see above). This was considered the “starting LECA node”. We rooted the phylogeny in the non-eukaryotic node farthest from the starting LECA node. The chosen rooting method was shown to be the most likely to not have a negative impact on the selection of the LECA group and its sister due to the lack of a proper outgroup (see **Supplementary Methods**). We next iterated through the sister clades of the starting LECA node and incorporated them into the LECA clade if:

1. The sister was mostly eukaryotic: that is the fraction of non-eukaryotic sequences was lower than or equal to 0.5, for sister clades with up to 20 sequences, or lower than or equal to 0.25 if the sister had more than 20 tips.
2. In the case that the first sister was mainly non-eukaryotic, but the second sister was mostly eukaryotic (according to the criteria above), the first sister was considered a transfer from the eukaryotic clade. In this case, we incorporated the first sister if all the following conditions were met:

a. the first non-eukaryotic sister had a number of sequences lower than 75% of the number of sequences in the LECA clade
b. the number of non-eukaryotic phyla was lower than 3 (assuming low diversity of the first non-eukaryotic sister), and
c. the fraction of non-eukaryotic sequences in the second sister is lower than 25%.

Once the LECA clade was defined with the above criteria, we removed it from the tree and performed the search iteratively until no more starting LECA nodes could be found in the tree.

For each LECA node, we selected only the eukaryotic sequences (mLECA-OG hereafter). Finally, we aligned these eukaryotic filtered sequences to compute a profile-based search of non-eukaryotic homologs using the above-mentioned criteria. We reconstructed the tree of the resulting set of eukaryotic and non-eukaryotic sequences using the phylogenetic reconstruction described above and identified LECA nodes in this tree following the same described procedure. This resulted in a final set of mLECA-OGs which is a refinement of the previous one. Only this final set was used in subsequent analyses.

mLECA-OGs were scanned for the possibility that they emerged due to HGT events between eukaryotic supergroups. For this, we located the LECA node in the phylogenetic tree, which by definition is ancestral to sequences from stem 1 and stem 2. We then inspected the two clades defined by the two first children nodes of the LECA node to see if either of the two nodes contained species from both stem 1 and stem 2, thereby containing the root of the eukaryotic tree. In 97% of the trees at least one of the children nodes satisfied this condition, making the LECA node unlikely the result of a HGT event between diverging supergroup taxa. Given that 29% of the LECA nodes are duplication nodes according to the species overlap algorithm, we searched for the first speciation node within the LECA node and repeated the analysis. In this case, 96% of the trees had children nodes that mapped to the root of eukaryotes, validating the results.

mLECA-OGs were also analyzed to assess the possibility of artificial splitting by OrthoFinder by clustering with MCL mLECA-OGs sharing at least 50% of their sister groups (see **Supplementary Methods**) which resulted in a minimal difference of our downstream inferences.

#### 2.6. Functional annotations of mLECA-OGs

We annotated all the proteins with KOfamScan v1.3.0 (https://github.com/takaram/kofam_scan)^92^ and HMM search against the COG database. Results from KofamScan were filtered in the following way: First, we selected those hits considered significant by KofamScan. When multiple hits were found, we selected the hit with the lowest e-value and highest hmmsearch score. In addition, for proteins with no hits passing the significance threshold of KOfamScan, we selected the hit with lowest e-value and highest hmmsearch score. To benchmark the accuracy of our predictions, we used 36 proteomes in our dataset that had been obtained from Uniprot and had mappings to genes in KEGG. This allowed us to establish KOs for a total of 154,248 proteins, independent of KofamScan. For those proteins we calculated the average correspondence of KO predictions per each species. We obtained 95% correct predictions on average, validating our KO annotation method. We further assigned each protein to a COG family by performing an hmmsearch using the COG profiles and selecting the first hit based on bitscore. We kept those COGs with a bit score higher than the percentile 90. Using the NCBI COG^93^ definitions table (https://ftp.ncbi.nlm.nih.gov/pub/COG/COG2024/data/cog-24.def.tab), we assigned the COG category to each protein.

Once all proteins were functionally annotated, we assigned a KO and a COG for each set of sequences (LECA-OGs, mLECA-OGs, or donor group, which in virus-mediated acquisitions would be the first prokaryotic sister) using the same approach used to functionally annotate eggNOG orthogroups. This approach takes into account the proportion of the KO and COG within the set of sequences and its overrepresentation with respect to the background KOs distribution (the whole set of annotated proteins). We chose a single KO and COG or multiple KOs and COGs in case of ties by minimising the sum of the KO rank in the proportions list and the overrepresentation list. To annotate the mLECA-OGs, we used the common KOs assigned to the LECA clade and its sister if they agreed, and the one of the LECA clade if they disagreed (see **Supplementary Methods** for details on the algorithm of assignation of KOs to a mLECA-OG). The mLECA-OGs with no non-eukaryotic homologs (innovations) with KOs just found in non-eukaryotic genomes were left unannotated. This can happen due to local homology between a non-eukaryotic KO and a eukaryotic protein.

#### 2.7. Inference of the LECA proteomes

For each eTOLDB version (A, B and C) and each LECA criterion (three supergroups and five supergroups), we obtained the set of mLECA-OG that fulfilled these criteria and inferred their origins and KO annotation. These proteomes are the ones that we used to infer the general proportion of acquisitions and innovations, as well as the general metabolism of LECA. Furthermore, we calculated a consensus proteome with the KO annotations that were pervasive and well-supported across the datasets to make inferences about the LECA features and metabolism. For this consensus we considered KOs present in two out of three TOLDB datasets or if at least five supergroups supported that KO in any TOLDB dataset.

#### 2.8. LECA metabolism and features inference

We inferred the presence of metabolic pathways and their completeness for each LECA proteome (one per eTOLDB version and LECA criterion) using the module anvi-estimate-metabolism of the Anvi’o v8^94^ package. We, finally, plotted the reconstructed metabolic map using ggKEGG v1.2.3^95^.

For other molecular components that are not necessarily associated to a single KEGG map such as the complexes that form the flagellum, the spliceosome or the cytoskeleton, we collected the set of genes associated to these components as depicted by KEGG brite, complemented them based on literature when necessary, and then checked the assigned origins of the different genes. We assessed the completeness of these features by extracting the subset of KOs that are general of eukaryotes (to minimise the effect of clade-specific KOs in the inference of this completeness) and checked the proportion of them present in our reconstruction of LECA by any database (for the origins) and the consensus proteome (for the presence and absence of the protein).

#### 2.9. Comparison with FLUPs, FLUOs and FLUAs

In order to benchmark the inferred LECA proteome, we compared it to the core and individual proteomes of selected unicellular, heterotrophic eukaryotic organisms. For this, we selected, from the eTOLDB proteomes, the P10K project^61^, and specific project repositories, organisms that were unicellular, heterotrophic free-living, and phagotrophs, autotrophs and osmotrophs (FLUPs, FLUAs and FLUOs, respectively, **Supplementary Table 5**). After cleaning and performing quality checks on the proteomes (see **Methods 1.1**), we functionally annotated the proteomes following the same protein annotation pipeline as for the eTOLDB genomes (see **Methods 2.5**).

Based on the KO predictions for each genome included in FLUPs, FLUOs and FLUAs we calculated the percentage of each genome that was shared with LECA in terms of number of KOs (number of shared KOs / number of KOs predicted in the eukaryotic proteome). We also obtained the frequency in which each COG functional category appeared in each genome within the three datasets as the number of genes that had a COG functional category divided by the number of genes with at least one COG category assigned.

### 3. Tracing mLECA-OG ancestries

#### 3.1. Identifying the origins of the mLECA-OGs

First, we analysed each tree by identifying the type of gene family among three categories: innovations, acquisitions and unknown. Innovations are those mLECA-OGs without non-eukaryotic homologs. We named acquisitions those having the LECA group nested within non-eukaryotic sequences. Among the acquisitions we identified two subcategories, the viral-mediated acquisitions, those pointing to a viral first sister followed by prokaryotic sisters, and direct acquisitions, those inferred to have acquired the gene directly from prokaryotes. We established an additional category for those trees whose directionality was not clear, the unknown origin mLECA-OGs. We classified trees as having an unknown origin when they had only viral homologs, less than 3 non-eukaryotic orders or fewer than 5 sequences. It is important to remark when considering these thresholds that the breadth of prokaryotic sequences has been heavily downsampled by the pangenomic approach. The sequences present in the sisters do not only represent themselves, but also a whole cluster of other prokaryotic sequences. Therefore each LECA clade may be deeper within the prokaryotic group and be nested while we only detect sister-to relationships.

To detect the non-eukaryotic groups that transferred a given gene family to the protoeukaryote, we analysed the first sister of the mLECA-OG in the case of the direct acquisitions and the first non-viral sister for the virus-mediated acquisitions. First, we discarded a non-eukaryotic origin for the mLECA-OGs for those trees where the sister clade contained more than 25% eukaryotic sequences, as although the sister would be mainly non-eukaryotic, the presence of eukaryotic sequences could be the result of secondary transfers from eukaryotes to a broad set of bacteria, hampering the identification of a confident sister. For the trees that passed this initial filter, we retrieved the proportion of each non-eukaryotic group present in the sister clade. Finally, to assign a taxon to the sister clade, we selected the most abundant group in the first sister (i.e., the one with the maximum proportion). In the case of more than one non-eukaryotic group with the maximum proportion in the first sister, we assumed that the clade was mixed and calculated its common ancestor, which could be Bacteria, Archaea, Viruses or a mixture of them.

Trees in which the first sister to the mLECA-OG was assigned to Nucleocytoviricota were assessed in more detail to assess the directionality of the transfer or the presence of secondary transfers from other groups. For this, we scanned the trees to search for additional, non-viral sequences. Given each tree with a sister in Nucleocytoviricota we located the point where LECA and its closest sister were located and went back in the tree towards the root in order to locate the first sister where the largest signal did not belong to Viruses. When such cases were found, the taxonomy of that sister was assessed and considered as the potential source of genes transferred to LECA through the viral lineage. Trees with only viruses as sisters and no other non-eukaryotic signal in the tree were classified as from unknown origin, as it was difficult to assess whether they were the result of a more recent HGT event from eukaryotes to viruses.

#### 3.2. “Stress test” for identifying the main non-eukaryotic contributors

To assess the congruence of LECA and its relationship to its sister, we checked the ultrafast bootstrap support (UfBS) values of the branch separating LECA and its first sister. However, the evolutionary depth of this project required a bootstrap support measure that accounted for broader taxonomic patterns rather than specific sequences moving through different nodes. Thus, we designed the taxonomic bootstrap, a measure of the support of a given taxonomic affiliation for a certain node of the tree. This version of the bootstrap calculates the proportion of ultrafast bootstrap trees that support the same taxonomic assignment for the sister of the ML tree regardless of the congruence of the sequences. In this case, we analysed the taxonomic bootstrap of the clade sister to LECA. To gain further insights about the assigned donor, we checked its proportion in the first sister and its presence in the second sister.

Given these measures, we used them in order to obtain the set of non-eukaryotic non-negligible donors beyond the assumed contributors (Alphaproteobacteria and Asgard archaea). We first established a set of parameters that were important for establishing reliable donors: the UfBS, the taxonomic bootstrap described above, the LECA criteria (3 to 9 supergroups) and the donor certainty (the proportion of the donor in the first sister). We then scanned the whole set of trees to obtain the thresholds for those parameters that maximised the proportion of trees assigned to Alphaproteobacteria and Asgard archaea, assuming that the selected thresholds properly remove noise and keep only the reliable signals. And then we checked which additional donors kept a significant number of trees using those same thresholds. Although the maximum Alphaproteobacteria and Asgard archaea proportion was given when using an UfBS threshold lower than 75%, we searched for the next maximum with at least 75% UfBS (**Supplementary Fig. 4**). The resulting thresholds that maximised the proportion of assignments to Alphaproteobacteria and Asgard archaea are: i) an UfBS higher than 75%, ii) a taxonomic bootstrap support higher than 95%, iii) seven supergroups criterion for the identification of LECA, and iv) a donor certainty higher than 95%. This resulted in the identification of 3 contributors beyond Alphaproteobacteria and Asgardarchaeota: 2 bacterial clades (Planctomycetota and Myxococcota) and a viral phylum (Nucleocytoviricota).

#### 3.3. Verticality test

To assess the possibility that bacterial signals emerged through vertical evolution of the Alphaproteobacterial donors we repeated the whole LECA prediction process as explained above. First we collected, out of the GTDB species representatives, a set of 35 representative Alphaproteobacterial proteomes. We selected a representative proteome per order (maximum CheckM completeness and, in case of ties, minimal CheckM contamination) and then randomly selected 35 orders out of this set. Then we filtered the pangenome to exclude all alphaproteobacterial protein sequences. Tree reconstruction and identification of ancestral Alphaproteobacterial groups were done as detailed above. An ancestral group was defined by the presence of five Alphaproteobacteria species, depending on the dataset. Then we identified the sister group to those ancestral groups and classified them into the different groups derived from the GTDB bacterial tree. So, for instance, S1 was represented by Gammaproteobacteria + Magnetococcia + Zetaproteobacteria. Then we counted the number of trees that had a sister in each of the categories (see **Supplementary Fig. 4b**). These values serve as the baseline vertical signal for each of the main donor groups, and show the expected proportion of trees that will appear associated to secondary donors rather than the main Alphaproteobacterial donor (see **Supplementary Fig. 4c**).

#### 3.4. Inferring the acquisition waves

We define a LECA gene acquisition wave as a set of genes potentially acquired from the same donor during the same period of time. To identify such acquisition waves, we computed the normalised stem lengths as in^4^ and plotted the distributions of these branch length values with respect to the taxonomic affiliation obtained according to the above procedure. For duplications occurring during eukaryogenesis and resulting in more than one LECA group, we considered the LECA node with the shortest branch length to the non-eukaryotic sister, as in^21^. This resulted in distributions with different modes for each taxonomic affiliation (**Supplementary Fig. 9**), whereas applying this approach to trees for OGs annotated to different functions resulted in overlapping distributions with similar modes (**Supplementary Fig. 10**). We inferred the acquisition module using the density function, the genes that are around the peak of the distribution are selected as members of the acquisition module. To obtain these genes, we set the threshold to the first minimum after the first maximum or, in its absence, the first inflexion point of the density curve (dashed lines in **Supplementary Fig. 9**). Those genes that transferred before the threshold (are in the peak) are included in the module. In contrast, the ones having longer stem lengths are discarded, assuming that they were transferred posteriorly or the values result from heterotachy or ghost lineages ^27,49^.

To infer the relative timing of the transfers and their timeline, we inferred the mode of the stem length distribution using a Bayesian framework^27^. In brief, this process uses a Markov Chain Monte Carlo algorithm to infer the posterior distribution for the parameters, mode and variance of a Gamma distribution, which has been shown to properly fit the empirical distribution of the stem lengths^27^. We used a pairwise comparison of the posterior distributions of the mode from different donors (assumed as relative ages) to establish a probabilistic timeline for the different transfers to the proto-eukaryote.

#### 3.5. Inferring the metabolism and features of the donors

To infer the ancestral features of the organisms that transferred genes to LECA, for a given donor group, we first selected, out of its genus representatives, the genomes sharing at least 50% of the genes transferred from that group to LECA in its acquisition wave, and annotated the genomes. These genomes constitute the “donor-like” genomes set. Second, we calculated the frequency of each KO present in these genomes. Then, we ran anvi’o v8^94^ with each donor-like genome. For both the metabolic modules and the KOs, we calculated their prevalence, that is the proportion of genomes containing a given KO or module. In the case of the metabolic modules, we transformed the prevalence by multiplying it to the mean module completeness in all the donor’s descendant genomes to take into account the completeness of the module. Using this metabolic reconstruction, we plotted the inferred donor’s metabolism using ggKEGG v1.2.3^95^. We further annotated each protein of the donor-like genome using an HMM search against the COG database and related the results with the COG categories. Using the annotated COG categories, we calculated the overrepresentations of each COG category proportion for a given donor with respect to the LECA consensus proteome. In this case, the overrepresentation against LECA consensus proteome would mean the functional enrichment of the donor’s contribution to LECA. Using the modules and the KOs prevalence, we assessed the presence and absence pattern of features relevant to the different eukaryogenesis models for all the donors. No statistical contrast for the enrichment was used as the sample sizes are non-homogeneous and, therefore, they do not provide enough statistical power, though we can study the general trends.

#### 3.6. Inferring Eukaryotic Signature Proteins (ESPs)

To define a new set of Eukaryotic Signature Proteins, as defined by^25^ we selected the KOs from the consensus proteome that were exclusively found as innovations.

## Acknowledgements

The authors want to thank members of the Gabaldón lab for the useful discussions, particularly Giacomo Mutti for his help in the taxonomy of the eukaryotic groups and Eduard Ocaña-Pallarés for the list of publicly available genomes of osmotrophs. We also want to thank Eugene Koonin and Victor Tobiasson for his comments on an earlier version of the manuscript. This project was mainly funded thanks to the support of the Gordon and Betty Moore Foundation (grant number GBMF9742). The authors thankfully acknowledge RES computational resources provided by BSC in MN5 to the projects BCV-2023-2-0019, BCV-2025-2-0018, BCV-2025-3-0020. TG group also acknowledges support from the Spanish Ministry of Science and Innovation (grant numbers PID2021-126067NB-I00, CPP2021-008552, PCI2022-135066-2, PLEC2023-010225, and PDC2022-133266-I00), cofounded by ERDF “A way of making Europe”, as well as support from the Catalan Research Agency (AGAUR) (grant number SGR01551); “La Caixa” foundation (grant number LCF/PR/HR21/00737), Fundació La Marató de TV3 (202328-31), AECC (PRYGN234923GABA), and Instituto de Salud Carlos III (IMPACT grant IMP/00019 and CIBERINFEC CB21/13/00061-ISCIII-SGEFI/ERDF).

## Authors contributions

MB, SM-M, and MM-H performed all bioinformatics analyses, wrote the necessary code, and produced the visualisations. TG conceptualised the project, obtained funding and supervised the project. All authors contributed to the design of the analytical steps, to the interpretation of the results, and to writing the manuscript.

## Competing interest declaration

The authors declare no competing interest.

## Data availability

All the alignment, trees, annotations, and supporting files to carry on this research are deposited in a Zenodo repository that will be available upon publication.

## Code availability

The code to carry on this research are deposited in a Zenodo repository that will be available upon publication.

## Extended Data legends

**Extended Data Fig. 1. COG categories distribution for free living unicellular eukaryotic genomes with different trophic strategies**. Each panel shows a trophic strategy, FLUA: autotrophs, FLUO: osmotrophs and FLUP: phagotrophs. Each point refers to a genome with a given trophic strategy and the black lollipop shows the proportion of that category in the consensus proteome of LECA.

**Extended Data Fig. 2. Acquisition waves in LECA**. Posterior distribution of the mode of the stem length distribution for each taxonomic affiliation. The panels show the results from different databases (in rows) and selection criteria (in columns).

**Extended Data Fig. 3. Probability of the waves occurring at different evolutionary relative time points**. Each heatmap value shows the pairwise probability of the wave from the donor in the row happening before the wave of the donor in the column. The donors are sorted by the acquisition time from older to more recent.

**Extended Data Fig. 4. Prevalence of pathways and enzymes in the genomes of the donor’s descendants**. Each box shows a group of metabolic pathways and enzymes. The opacity of the colour shows the prevalence (the percentage of genomes with the feature) of that specific pathway or enzyme in extant genomes from the donor’s clades that share at least 50% of the KOs that these donors transferred to LECA. Abbreviations, AA degr.: amino acid degradation; Aut.: autotrophy, Ferm.: fermentation; Hase: hydrogenases; Het.: heterotrophy; N metabolism: nitrogen metabolism; OXPHOS: oxidative phosphorylation; Phot.: photosynthesis; S metabolism: sulphur metabolism.

**Extended Data Fig. 5. Origins of the pathways and enzymes in the consensus proteome of LECA**. Each box shows a group of metabolic pathways and enzymes. The opacity of the colour shows the origin’s proportion. For each KO, we calculated the proportion of origins for all the databases assigned to a clade, and weighted to the total number of KOs in the module, resulting in a proportion for the feature. Abbreviations, AA degr.: amino acid degradation; Aut.: autotrophy, Ferm.: fermentation; Het.: heterotrophy; OXPHOS: oxidative phosphorylation; Phot.: photosynthesis; S metabolism: sulphur metabolism.

## Notes

### Competing Interest Statement

The authors have declared no competing interest.

### Summary of Updates

Some analyses were repeated based on revision by peers

