## Supplementary text and figures for "Diverse gene ancestries reveal multiple microbial associations during eukaryogenesis"

### Supplementary Information for: Diverse gene ancestries reveal multiple microbial associations during eukaryogenesis

#### Supplementary Methods

##### ***Phylogenetic backbone and supergroups for eukarya***

In order to determine the presence of a given gene in LECA a certain distribution of species was required. In order to establish clear patterns we distributed species into two stems and nine supergroups. As a ground phylogeny for the analysis we chose a consensus between four topologies Lax et al.[1], Tikhonenkov et al.[2], Torruella et al.[3] (for the internal topology of the eukaryotic tree), and Williamson et al.[4] (to place the root of the tree). The two stems were separated in Diphoda+: Diphoda + Provora, *Meteora* and Hemimastigophora (PHM) + Discoba (Stem 1), and Opimoda+: Opimoda + Metamonada (Stem 2). This stem is also supported by the last attempts on rooting eukaryotic phylogeny[4]. Within each of the stems, a set of Supergroups was defined. Starting within Diphoda+ (**Supplementary Fig. 1**), the three reference trees recover the monophyly of Cryptista and Archaeplastida, therefore, we considered both of them as an artificial supergroup named Archaeplastida plus. Picozoa was included in archaeplastida as sister to Rhodelphidia and Rhodophyta as shown by Torruella et al.[3] and Schön et al.[5]. The monophyly of SAR is well established in the literature, however, the position of telonemids and haptists is still debated[6], although they are always closely related to SAR. The three trees recover the monophyly of these groups, although they do not resolve their internal topology, thus, we considered them a polytomy and grouped them into another artificial supergroup that we named TRASH just for labelling purposes in our analysis. Provora is a recently discovered eukaryotic group which was originally placed within Diaphoretickes[2]. Recently, with the sequencing of *Meteora sporadica*[3,7,8] and Hemimastigophora species[1], these three groups have formed a monophyletic group with good support[3] which we have considered as a supergroup and have named PHM. Discoba appears as sister to all Diaphoretickes in Tikhonekov et al.[2], as sister to PHM and Diaphoretickes in Torruella et al.[3] and as a polytomy in the root of eukaryotes in Lax et al.[1]. Therefore, we put a polytomy in the early diversification of Diphoda.

The other considered stem is the artificial group containing Opimoda+: Amorphea, CRuMs, Malawimonadida and Ancyromonadida[9], which constitute Opimoda, and Metamonada. The three topologies agree in the monophyly of the Amorphea clade, which is composed of Amoebozoa as sister to Obazoa (Breviatea, Apusomonadida and Opisthokonta). CRuMs appear as sister to Amorphea in the three topologies. Malawimonadida and Ancyromonadida are the most conflicting nodes between these three topologies. In Lax et al.[1] and Tikhonenkov et al.[2] Ancyromonadida appears as a sister to Metamonada. However, with the increased number of new transcriptomes in Torruella et al.[3], these groups form a monophyletic group sister to Amorphea and CRuMs (the clade Podiata[10,11]) with full support with ML, 0.99 with BI (CAT+GTR) and 0.75 for BI with (CAT-Poisson) model. According to this, we decided to place Malawimonadida and Ancyromonadida in the Opimoda group, although we considered them polytomous and two independent supergroups in our analyses. Finally, Metamonada was artificially considered in this stem to balance the tree. This clade, with many parasitic representatives, forms a monophyletic group in the three topologies, although it cannot be properly placed as sister to any other eukaryotic group. Therefore, we considered it as another supergroup for our analyses. The internal topology of Metamonada has been determined according to Stairs et al.[12] and Eglit et al.[13]

##### *Rooting method*

The rooting of the gene trees is an important aspect of a phylogenetic based analysis. Whenever possible, outgroups should be added to ensure that the root is correctly placed. In an analysis like the one presented here, the rooting placement cannot be based on outgroups as there are no known ones and therefore we need to use approaches that will not affect the determination of the LECA group and its sister while not necessarily placing the root in the correct position. In order to do this, we decided to place the root at the non-eukaryotic, most distant possible branch from the initially detected LECA node based on branch lengths.

To assess the rooting method and ensure our LECA prediction and its sister was as correct as possible, we tested this approach in a set of trees containing 78 species with a known taxonomic relation among them. The trees are part of the Quest for Orthologs[14] initiative and are based on a set of carefully selected and curated genomes that contain both eukaryotes and prokaryotes. These trees are reconstructed using a species as a starting point (seed species) and each tree has a known sequence that was used to run the initial homology search that then resulted in the reconstructed tree. We selected the trees reconstructed for *Homo sapiens*, *Saccharomyces cerevisiae* and *Arabidopsis thaliana*. The aim of this analysis was to see how the rooting method affected the prediction of the sister group to Metazoa (*H. sapiens* trees), Fungi (*S. cerevisiae* trees) and Plants (*A. thaliana* trees). We first rooted the trees based on the known taxonomic relationship between species, placing the root at the species farthest related from our seed species and among those we selected the sequence that had a longer branch length (get_farthest_oldest_leaf() method from the ETE3[15] package). We then selected the monophyletic group of interest containing the seed sequence and checked its sister. This process was repeated twice more using two different rooting methods: midpoint rooting and the rooting approach used in the LECA analysis (placing the root in the farthest non-metazoan, non-fungal and non-plant node).

When compared to the taxonomic based rooting, the midpoint rooting is only able to recover the same sister species in 69% of the cases for Metazoa, 73% for Fungi and 75% for plants. In contrast, the LECA approach is able to recover the sister correctly in 89% of the cases for Metazoa, 88% for Fungi and 90% for plants. These results point to the fact that, despite lacking a proper outgroup where to root our gene trees, we are likely to obtain a reliable LECA and its sister determination in our analyses.

##### *Functional annotation of the mLECA-OGs*

To annotate the mLECA-OGs following the procedure described in Methods, we separately obtained an annotation of the LECA and the sister groups. For each group (prokaryotes, eukaryotes and viruses), we used a different subset of the available KOs (**Supplementary Fig. 11**), those specific to each group. We also retrieved the intersections, when a KO was present in combinations of these groups (**Supplementary Fig. 11**, left panel).

For the sister annotation, we considered all the KOs present in prokaryotes and viruses, regardless of their taxonomic or functional deepness, assuming that the donor could be specific. Whereas in the case of the LECA, where we need to be general, we annotated the clade using functionally and taxonomically general KOs. We excluded from the algorithm KOs that mapped only to specific eukaryotic groups, such as, fungi or animals (according to the KEGG organisms taxonomy) and KOs exclusively with functions associated to very specific functions, such as cancer- or infection-related KOs. In case of a KO present in both a specific and a general functional pathway (e.g., a KO present in the colorectal cancer set of KOs but also present in the glycolysis), we kept it.

Once these KOs were filtered from the list of KOs in the LECA and the sister groups, we retrieved the annotation of the mLECA-OG as follows: if there were common KOs between LECA and the sister groups, we kept the common KOs as the annotation for the mLECA-OG. In the absence of common KOs, if the LECA filtered KO was available, we assigned it to the full orthogroup. When the LECA KO was missing due to filtering, we assigned the KO of the sister group. Otherwise, the mLECA-OG was not annotated. This process allowed us to properly annotate innovations with eukaryotic KOs, removing possible artifacts caused by local homology to non-eukaryotic KOs. Using this annotation approach we covered 98.29% of the mLECA-OGs on average. The remaining non-annotated proteins mostly belong to innovations and unknown origins mlECA-OGs.

##### *Robustness of donor metabolic feature inference*

Due to the inherent bias towards better-sampled clades within databases (**Supplementary Fig. 12)**, we implemented a re-sampling with replacement strategy to assess the dependency of the inferred metabolic potential to the taxonomic representation of the database. For this, we first selected one representative per genus, and kept the genomes sharing at least 50% of the genes transferred from that group to LECA in its acquisition wave, and annotated these genomes (967 Alphaproteobacteria, 12 Heimdallarchaeia, 60 Asgardarchaeota, 67 Myxococcota, and 144 Planctomycetota genomes). We then selected 100/n genomes per class (or order, for Alphaproteobacteria and Heimdallarchaeia classes), n being the number of classes (or order), and rounding decimals to the nearest whole number. For classes that had less than 100/n genomes, one genome may have been chosen more than once, to reach the quota. This process was carried out 100 times, and we calculated the prevalence of each KO and modules across all iterations (**Supplementary Fig. 13**).

#### Supplementary Discussion

##### *Pre-LECA duplications not inferred by the orthology calling*

To assess the impact of LECA gene families that are split by OrthoFinder[16] during the orthogroups inference, we joined mLECA-OGs by sister concordance. Those mLECA-OGs sharing at least 50% of their sister groups were clustered with MCL using the sisters' overlap value as weight. This resulted in clusters of OrthoFinder OGs that were submitted to the search of non-eukaryotic sequences and the tree reconstruction pipeline (detailed in Methods). This procedure resulted in three different types of mLECA-OGs: i) separated as independent mLECA-OGs, ii) pre-LECA duplications, and iii) LECA clades within an already inferred mLECA-OG that may result from transfer, duplication and loss dynamics.

We analysed the impact of this clustering by re-computing general statistics of the LECA proteome sizes and donor assignation. For all the databases, the proteome size decreased 5% (around 500 families were joined or lost as they failed the LECA criteria). Regarding the impact of this reduction in the assignment of the donors, we re-ran the donors' detection algorithm and the results were the same for the joined and initial tree datasets. We further calculated the percentage of each donor for each proteome using the joined and the initial tree datasets, obtaining a maximum difference of 0.37% among the prokaryotic contributors and 1.29% difference for Nucleocytoviricota.

***Selection of putative donors using the "stress test"***

To define the criteria used to define a potential donor to LECA, we assessed the effect of varying the stringency of thresholds on three parameters that impact the certainty about the taxonomic assignment of the sister-clade to the mLECA-OG (the ultrafast bootstrap of LECA and the sister, the taxonomic bootstrap and the proportion of the assigned donor in the first sister). When varying these thresholds, we compared the fraction of sister-clade assignments corresponding to the two already-known partners in LECA (Asgardarchaeota and Alphaproteobacteria) among the trees that passed the thresholds. The rationale of this approach is that assignments to these known partners are likely to be true calls, and the fraction of false calls are expected to decrease as more reliable trees are selected with more stringent thresholds. Therefore assessing the relative number of Asgardarchaeota and Alphaproteobacteria with respect to other assignments can be informative to define which thresholds maximise the selection of LECA genes that are sisters to these groups.

The distributions of ultrafast bootstrap support (UfBS)[17] values in both the LECA (64.8% of the trees have a UfBS higher than 75) and the LECA-sister branches (41.3% of the trees have a UfBS higher than 75) show that most of the nodes of interest are supported with an UfBS > 75% (**Supplementary Fig. 14a,b**). Regarding the congruence of the LECA group across the UfBS trees, 71.2% of the trees have a congruence higher than 95%, showing that the LECA definition is robust across the different UfBS trees (**Supplementary Fig. 14c**). The taxonomic bootstrap distribution is also left-skewed, meaning that the values are concentrated in the upper values, supporting a robust donor assignment (**Supplementary Fig. 14d**).

We assessed three levels of stringency (low, medium and high) of three support variables (UfBS, taxonomic bootstrap, and donor support) and the number of supergroups to assess the relative contribution of Asgardarchaeota and Alphaproteobacteria to LECA (**Supplementary Fig. 4**). The most influential factor was the number of supergroups used to define a mLECA-OG. Increasing the number of required supergroups leads to significant increases in the relative fraction of trees with Asgard archaea as sister groups while the proportion of those traces to Alphaproteobacteria diminishes. The other stringency factors reduced the total number of trees selected while having a small impact on the relative proportions of Asgard archaea and Alphaproteobacteria, suggesting that different taxonomic assignments of the donor groups have similar phylogenetic support but may differ in their extant representations across eukaryotic supergroups. These results are compatible with the major contribution to core housekeeping functions of Asgard archaea and to metabolic functions of

Alphaproteobacteria and other donors, as metabolic genes are more likely to be lost in specific lineages.

To select a minimal set of potential donors with reliable contributions, we established our thresholds in conditions that maximise both the relative proportion of Asgard archaea and Alphaproteobacterial contributions, and the number of considered trees. These thresholds were: medium UfBS of the branch connecting the LECA group with its sister (>= 75%), high taxonomic bootstrap (>= 95%), high donor proportion (>= 75%) in both first and second sister clades. We selected donors contributing more than 20 genes passing these thresholds in all eTOLDB datasets. The selected major donors were five non-eukaryotic groups: Nucleocytoviricota, Asgardarchaeota, Alphaproteobacteria, Planctomycetota and Myxococcota.

***Rationale of the verticality test***

The topology of gene phylogenies capture the evolutionary processes that shape the evolution of genes, including gene acquisition, gene loss, gene duplication and HGT. The aim of this study is to assess the role of HGT during eukaryogenesis, and define which non-eukaryotic taxonomic groups contributed to the earliest eukaryotic lineages. Trying to track such ancient HGT in gene phylogenies reconstructed with extant sequences is challenging, as many subsequent events may have muddled the signal. In particular, when finding that two closely related taxonomic groups are HGT donors, an obvious question to ask is whether the two groups were independent contributors or whether subsequent events such as gene loss or inter-domain HGT could have shifted the signal resulting in two similar signals from a single true donor. In this analysis, we have found evidence that the main recognised donors (Alphaproteobacteria and Asgard archaea) played a role in eukaryogenesis, as expected. But we also find additional signals from other bacterial groups.

The verticality test applied here is an attempt to distinguish whether those additional donors are independent donations of genes to LECA or they are the result of technical (poor sampling) or alternative evolutionary events (gene loss). The test consists of determining a baseline of what we can consider vertical evolution in the face of such events, or said in a different way, in which proportion can we expect alternative sisters to appear if closer homologs are not present the main donor.

In order to assess this, we built a new dataset containing 35 Alphaproteobacteria (**see Methods**). Then we removed the species of the main donor from the pangenome. Then, we run the dataset through our tree reconstruction pipeline. We modified the LECA detection pipeline to capture groups of ancient Alphaproteobacteria genes (see **Methods**) and defined the donor group with the same algorithm we use for LECA. The expectation is that most of the trees will have a sister in the closest taxonomic group as defined by the GTDB species tree (called S1, for Sister 1). So, in this case, we expect a main signal in Gammaproteobacteria, Magnetococcia and Zetaproteobacteria which in this analysis represent the closest sister clade to Alphaproteobacteria. But there is going to be a fraction of genes that will have lost this S1 homolog and will map to other taxonomic clades (named S2,

S3, S4 and so forth). For a purely vertical evolution the expectation is that the farther away the sister is the fewer trees will map to it. And this is exactly what we observed in this test.

In **Supplementary Fig. 4c**, we can observe a descending signal corresponding to farther related sisters. If the signals observed in the LECA analysis were the result of purely vertical evolution derived from the Alphaproteobacterial donors, we would expect the trees to show similar patterns. Instead we find strong signals in three sister groups (S3 which contains Myxococcota and S7 which contains Planctomycetota and finally S6 which contains many of the groups that provided weaker signals such as Chloroflexota or Firmicutes). The strong signals found in these three distant sister groups deviate from the pattern formed by vertical evolution, leading to the conclusion that non-vertical evolutionary processes, such as HGT, underlie these signals.

***Metabolic reconstruction and the taxonomic breadth of the inferred modules***

Following the strategy of the most recent common ancestor (MRCA) calculation for the KEGG orthologs (KOs), we identified MRCAs for each functional module. Therefore, we were able to select those that whose MRCA was LUCA or eukaryotes, excluding those modules found only in a narrow group of eukaryotes (e.g., animals, plants, etc.) or functionally restricted (e.g., cancer, infectious diseases, etc.). This resulted in a less complete metabolic map (**Fig. 1**) as compared to the metabolic map with all the complete pathways found using our annotation (**Supplementary Fig. 15**). However, although the full map is more complete, we observe the basic metabolic features displayed in all eukaryotes, such as the central carbon metabolism, some components of the ATP synthesis, basic biomolecules and cofactor synthesis pathways.

Among the manually discarded modules, we find Reductive pentose phosphate cycle (Calvin cycle) (M00165), CAM (Crassulacean acid metabolism) in light (M00169), and C4-dicarboxylic acid cycle, phosphoenolpyruvate carboxykinase type (M00170), which are related to carbon fixation. Moreover, the eukaryotic ATPase (V-type) (M00160) is only annotated in animals in KEGG, showing the existing bias towards annotations based on model organisms. We manually included it in the final reconstruction as it was complete (**Supplementary Table 4**). The maps in KEGG are based on extant genomes projected on known pathways for model organisms. However, LECA lived ~1.5 - 2.5 Ga, resembling more non-model unicellular organisms, and the cellular mechanisms may have substantially changed during this period. Therefore, the more general the annotated modules, the more reliable our reconstruction is, although the filtered annotation of modules is still sensitive to the changes that happened during the evolutionary history of eukaryotes. Based on these premises, we decided to use the filtered set of general modules in all downstream analyses.

***LECA reconstruction***

*Limitations of automated reconstructions*

Reconstruction of the repertoire of proteins likely present in LECA is a necessary step to gain insights into the phylogenetic ancestries of LECA –the main purpose of this study. For this, we used an automated approach following procedures that are state-of-the-art in the field.

Although more sophisticated reconstructions benefiting from the manual curation of expert biochemists have been performed for some pathways and cellular structures (reviewed in refs.[18–21]), they do not provide a full picture of the LECA gene repertoire and do not include the most recent genomic datasets. Automated reconstructions of LECA, such as the one performed here, have been used in previous studies inquiring about the nature of LECA or other ancestral organisms[20,22–25] and we consider it appropriate for the aims of this study. Nevertheless, the limitations of such an automated procedure should be acknowledged in particular with respect to the functional annotations of LECA families, which are necessarily based on annotations of extant proteins available in the mined databases. Limitations of family-based orthology approaches –such as the eggNOG strategy used here for functional annotation– have been extensively discussed, in particular in difficult cases due to domain shuffling or low sequence similarity[26,27]. We thus stress that the purpose of this reconstruction is to approximate gene families and functional categories likely present in the ancestral LECA proteome, rather than to provide a carefully curated and detailed picture of LECA's traits. The comparison of our automatically reconstructed LECA proteome and its functional annotations with earlier work, including some expert-curated datasets, serves to assess the completeness and validity of our reconstruction.

To infer the cellular mechanisms and structures that were likely present in LECA, we examined annotations in the consensus proteome (**Supplementary Data**), which is the set of KEGG orthology terms (KOs) that were present in at least two eTOLDB versions or supported by a LECA criterion of five supergroups. Note that there is not an exact correspondence between KOs, which we use to approximate functional annotations to OGs, and the OGs reconstructed here. This means that an OG may be restricted to eukaryotes, but its functional annotation is described by a KO that has a broader distribution, perhaps due to domain sharing with other OGs. This also means that some of our functional annotations are broad and may not specifically describe the specific function within eukaryotes. Despite these limitations, KO-based annotations provide a useful means to explore the metabolic completeness and the overall cellular features likely present in LECA. To minimize annotation artefacts and keep a broad functional characterisation, we used a taxonomy-informed approach. We annotated eukaryotic proteins with taxonomically and functionally broad eukaryotic KOs and KOs present in both prokaryotes and eukaryotes (the LUCA set of KOs, **Supplementary Fig. 11**, see **Supplementary Methods**). Similarly, we annotated prokaryotic sequences with KOs that were only found in prokaryotes or in the LUCA set. The reconstruction of the cellular features of LECA (**Supplementary Table 7**) agree with the current consensus that LECA was a complex organism showing most of the features present in extant free-living unicellular eukaryotes[19]. To expand the description provided in the main text, we here discuss some relevant observations in different key biological processes and cellular structures. By highlighting congruent findings with previous detailed studies, we underscore the validity of our reconstruction.

*DNA and RNA processing*

Our LECA inference depicts almost complete modern DNA and RNA processing systems (e.g., polymerases, transcription factors, ribosome, etc.), something that has been observed in previous studies[21,28–30]. Similar to previous more detailed analysis[31], we reconstruct a complex LECA spliceosomal machinery, where all its components are present. As our reconstruction focuses on proteins, we are missing the snRNAs that are part of the spliceosome, but, given that the remaining components are present, we expect those snRNAs to also have been present in LECA. In a previous study, Vosseberg et al. performed a reconstruction of the LECA spliceosome[31], eight of the proteins were deemed possibly part of LECA. We searched for these proteins in our reconstruction and were able to find PPIE (K09564), PPWD1 (K12736), PPIL3 (K12734), PRPF38B (K12850), ZNF830 (K13104) and DDX42 (K12835). We only missed two of the proteins: SFPQ (K13219) and PNN (K13114), although this last one appeared as a potential innovation in the TOLDBB –3 supergroups reconstruction. Many of the proteins detected as part of the spliceosome are classified as innovations (33%). Still, other proteins have putative bacterial origins (37%) (see **Supplementary Table 7**).

The LECA ribosome has almost all the proteins of the current eukaryotic ribosomes, only missing RP-L41e (RP-L41e), with a dominating archaeal origin (52% of the ribosomal proteins) (see **Supplementary Table 7**). In a previous study large proportions of genes related to ribosome biogenesis were identified as having emerged in LECA such as NGDN, PARN, RPP25, etc. and are also identified in our analysis[32]. In fact, most of the complexes described in KEGG pertaining to Ribosome biogenesis are complete in our reconstruction of LECA, the only exception being the t-UTP complex which is missing UTP8 (K14547) and UTP9 (K14551). These missing proteins are likely undetected due to the applied KO annotation filtering regarding KOs present only in few eukaryotic species, as both proteins appear associated only to Fungal species in KEGG. Additionally, due to their limited distribution, it is likely that they were indeed not in LECA.

The RNA degradation system is inferred to be complex in LECA. It includes, for instance, the eukaryotic core exosome, the decapping complex and the Pan complex which are all complete in our LECA reconstruction. Other specific proteins associated to the mRNA degradation system (NMD3p (K07562), UPF1 (K14326)) and the poly(A)-degrading complex (POP2 (K12581) or CCR4 (K12603)) are also recovered in our LECA reconstruction[33]. Some of the proteins seem to be missing, which in some cases can be attributed to not being in LECA (e.g., SKI7 (K12595) and CAF120 (K12609) are only found in fungal species). The main signals corresponding to the RNA degradation system are split between innovations (25%) and bacterial origin (29%) (see **Supplementary Table 7**).

We infer nearly complete DNA replication, mismatch repair and non-homologous end-joining mechanisms, base excision repair and nucleotide excision repair pathways. We recover the complete DNA replication pathway and  the mismatch repair pathway. Homologous recombination pathway is missing the DNA endonuclease RBBP8 (K20773), and proteins that are species specific in animals or yeasts such as (BRCA-1, Abraxas, or XRS2). Both DNA replication and homologous recombination had a main contribution of archaea (31% and 21%, respectively) and non-negligible bacterial contribution to both systems (14% and 25% of the genes, respectively) (see **Supplementary Table 7**).

The polymerase regulation that we observe in extant eukaryotes is also present in LECA, although the origin of the proteins involved in this mechanism is uncertain. The common and core subunits of Pol I, Pol II and Pol III were present in our LECA reconstruction as were most of the Pol specific units, only missing subunits A14 (K03001) and A34 (K03003) of Pol I, both having been associated in KEGG with only fungal species and as such would not appear in LECA in our reconstruction. These and other genes involved were already described as part of LECA[34].

The nuclear pore complex was previously described as having been present in LECA [35,36]. We infer the basic components of the nuclear pore complex (NPC) as part of our LECA reconstruction: cytoplasmic fibrils, cytoplasmic ring, central channel, spoke complex, lumenal ring and nuclear basket. Some proteins of these different complexes are missing, for instance the basket lacks 3 out of 7 proteins (Nup2, Nup153 and Nup60, though this last one appears to be specific for fungal species and as such should not be part of LECA[37]). The nuclear transport complexes show that LECA's NPC allowed the export and import of localisation and export signals. We only appear to miss the PHAX protein (K14291) which is a RNA export protein. We infer a complete transcription export complex and an exon junction complex nearly complete (15/17 - we miss MLN51 (K14323) and Pinin (K13114), the last one appearing in TOLDBB as an innovation). Based on our analysis we assess that 45% of the proteins of the nuclear pore complex were inferred to have evolved during eukaryogenesis, as we detect them as innovations. The following most abundant donor is bacteria, inferred to have transferred 26% of the genes of the nuclear pore complex (see **Supplementary Table 7**). Further detailed analyses on these proteins may shed light on the evolution of the nucleus.

*Protein processing*

Protein processing was already complex and similar to that of extant eukaryotes with functional endoplasmic reticulum (ER) and Golgi apparatus, which could modify, export and degrade proteins[21,38,39]. We detected a complete Sec-dependent translocation pathway, including the eukaryotic channel proteins, a complete signal recognition particle (SRP) and SRP receptors. We also detected signal-dependent and -independent channel proteins, allowing LECA to export proteins in different ways. These mechanisms mainly evolved through innovation during eukaryogenesis, although with a non-negligible signal from bacteria (28%), with the detected specific contributions being from  Asgard archaea (5%) and Planctomycetota (4%) (see **Supplementary Table 7**).

We further investigated the ER processing system of the LECA proteome, which contained the essential protein processing functions observed in current eukaryotes (translocons to import the protein coupled to the ribosome, proteins that detect misfolding and target them to be degraded by the proteasome). These results suggest that the ER evolved during eukaryogenesis until its almost current complexity in LECA. The degradation of the proteins was similar to that of extant eukaryotes, as LECA had most of the complexes required to perform the ubiquitin-mediated proteolysis and the core functional proteins of the proteasome. The different mechanisms evolved mainly in the eukaryotic stem lineage and from Bacteria. However, there are bacterial and archaeal contributions in all the mechanisms. For instance, the proteasome shows 36% of the proteins having archaeal origins of which 17% are from Asgard Archaea. This is compatible with smaller-scale studies such as that of the ubiquitin signalling system[40]. Finally, some genes are also inferred to have been transferred through viruses (16% of the proteolysis system) (see **Supplementary Table 7**).

*Transport and catabolism*

Extant eukaryotes transport proteins and other cellular molecules by intracellular vesicles; the machinery of this kind of transport was present in LECA. With our reconstruction, we infer a Golgi apparatus with its essential proteins and stable communication with the ER by the Stx18 (K08492), Use1 (K08507), Syp7 (K8506) and Sec20 (K08497) genes. Another mechanism of transport from the extracellular media is endocytosis. Endocytosis was previously described as being mostly present in LECA, with genes such as clathrin (K04644), epsin (K12471) or eps15R (K12472) being largely distributed in different species[21,41]. According to the presence of these genes, LECA had an endocytic system similar to many currently eukaryotic taxa which was later expanded in some species such as *Saccharomyces cerevisiae* or *Trypanosoma brucei*. Our reconstruction of LECA also recovers a complex endocytic system, and includes the ESCRT systems, as well as other proteins that facilitate this process in both clathrin-dependent and -independent cases. LECA also had the two different stages of the endosome (early and late), showing that these mechanisms evolved during the eukaryogenesis from earlier and more simple vesicle systems. We finally infer that LECA had a limited number of ABC transporters, we have only found three members of the ABCA subfamily (1, 3 and 5), five from the ABCB subfamily (1, 6, 7, 9 and 10), four of the ABCC subfamily (1, 2, 4 and 10), three of the ABCD subfamily (3, 4 and PXA1/2) and two from the ABCG subfamily (ABCG2 and PDR5). Many of these transporters may have duplicated after the diversification of eukaryotes[18,42–44]. The origin of this system is mixed, with innovations and proteins that originated in bacteria and were transferred to the proto-eukaryote or already present in the bacterial partner of the proto-eukaryote. In this system, the contribution of archaeal donors is negligible (~3%) except for the endocytosis, in which we detect around 19% of archaeal contribution (specifically in the ESCRT system, concordant with previous assessments[45]) and strong evolution of the system during eukaryogenesis as 35% of the proteins are inferred to be innovations (see **Supplementary Table 7**).

Once outer bodies are incorporated, the phagosome and the lysosome process them. The ability of LECA to phagocyte other organisms is a heavily debated topic. With some authors arguing that LECA incorporated the alphaproteobacterial genome through phagocytosis and others indicating that this ability came later and that the incorporation of the alphaproteobacterial species was accomplished through symbiosis[44]. Our reconstruction of LECA contained numerous proteins from the phagosome, though a large proportion of the missing genes have only been described in animals and therefore do not appear in our reconstruction. Among the detected proteins we find the scavenger receptor SRB1 (K13885), a nearly complete phagocytosis system including VAMP3 (K13505), Stx13 (K13813), Rab5 (K07888) and Rab7 (K07897), F-actin (K05692), VPS34 (K00914), Dynein (K10413 to

K10416), TubA (K07374) and TubB (K07375), Hrs (K12182), the full vATPase complex (K02144 to K02155), cathepsin (K01365) and Sec61 (K10956, K09481, K07342), though we are missing some proteins such as MPO (K10789) which catalyzes the conversion of $H_2O_2$ into a potent antimicrobial agent or EEA1 (K12478) that plays an important role during the early stages of phagosome maturation. The lysosome we can infer in LECA was able to regulate the ATPase V-mediated acidification, as it contained a nearly complete ATPeV complex and its regulators DMXL (K24155) and WDR7 (K24738). We also infer that it had most of the lysosomal membrane proteins such as LIMP (K12384), NPC (K12385 and K13443) or cystinosin (K12386). We may be missing some of the minor lysosomal membrane proteins that are more widely distributed in Eukaryotes such as sialin (K12301) and CLN5 (K12390). Regarding the function of the lysosome, we identify all the general eukaryotic proteases except the napsin (K08565), 11 out of 13 glycosidases (missing HYAL1 - K01197 and GALC - K01202), lipases, etc. This suggests LECA could process outer bodies through the phagosome and the lysosome. The KOs of the phagosome have mixed origins, our reconstruction suggests that 25% of this structure may have evolved through eukaryotic innovations, 20% acquired from bacteria, 29% from archaea, with 6% of the phagosomal proteome detected to have been transferred by viruses. The lysosome follows a similar pattern with 31% having been obtained from innovations, 30% from bacteria, 13% from archaea and only 1% having been obtained through virus-mediated acquisition (see **Supplementary Table 7**).

Peroxisomes are involved in oxidative reactions such as lipid catabolism. Similar to previous, more detailed studies (reviewed in[46]), we found that LECA has a complete membrane importer and matrix protein import systems. The main peroxisomal functions (fatty acid oxidation, phospholipid biosynthesis, sterol precursor biosynthesis, amino acid, antioxidant, and purine metabolisms) are present in the reconstruction of the LECA peroxisome. According to our reconstruction, the peroxisomal proteome has mostly bacterial origins (50%, 14% from alphaproteobacterial origins and 6% from Myxococcota) with a significant share of innovations (20%) (see **Supplementary Table 7**).

*Cell cycle*

Eukaryotic cells need to synchronise several processes to start a division as their cellular complexity is higher than prokaryotes, this is controlled by the cell cycle. In previous studies it has been observed that LECA had a complex cell division machinery[21,47,48]. We confirm this observation with our reconstruction of LECA, where we find several proteins involved in each step of the process. Studies in eukaryotic model organisms such as Human, Yeast and the algae *Chlamydomonas reinhardtii* have shown that genes involved in cell cycle can vary greatly among organisms but that a core set of genes are maintained across different clades and are assumed to have been present in LECA[49,50]. Our LECA model contains most of the genes that are predicted as shared between Human and the algae *C. reinhardtii*. For instance we find the cyclin dependent kinases CDK1 (K02087), CDK2 (K02206) and CDK7 (K02202), some of the cyclins such as CycA (K06627), CycD (K10151) and CycH (K06634), or nearly all proteins from the Anaphase promoting complex/cyclosome (APC1 - K03348 to APC12 - K03359, APC13 - K12456 but missing APC15 - K25228 and APC16 -

K25229 that have only been found in animals in the KEGG database). A PLK1, which is an important and highly conserved mitotic regulator, is found in 5 copies in humans but only one in yeasts and *C. reinhardtii*[49]. While this could indicate that the five copies in the human genome emerged through duplications we find that our model of LECA contains two copies of the gene corresponding to PLK1 (K06631), which is involved in the cell cycle, and PLK4 (K08863), which is involved in the Forkhead box O (FOXO) signalling pathway which is involved in apoptosis. Genes belonging to the cell cycle in our LECA reconstruction were obtained through a combination of archaeal and bacterial acquisitions (17% and 22% respectively) and innovations (28%) (see **Supplementary Table 7**).

The evolution of eukaryotic sex is also a complex question that requires cellular studies. We found that LECA could have carried out meiosis, as its proteome contained proteins in all the main subprocesses of the meiosis, as supported by earlier detailed studies[51–53]. However, as observed in the cell cycle, the regulatory network of the meiosis was still not constituted. The origin of meiotic genes in LECA are scattered, most of them are innovations during eukaryogenesis (24%) and the rest are mostly archaeal (22%), bacterial (22%) and some are inferred to have been transferred through viruses (9%) (see **Supplementary Table 7**).

Autophagy allows some cells to survive by obtaining resources from the degradation of subcellular structures. The consensus proteome of LECA indicated that it had a modern complete autophagy process, similar to what had been previously inferred[54]. We also infer that LECA was able to induce a basic process of apoptosis (induced cellular death), as we found genes in specific positions of the different stages of the apoptosis process, such as DNA fragmentation (ENDOG - K01173 and AIF - K04727). Genes related to other variants of induced death (necroptosis and ferroptosis) are also inferred in LECA. These features are mainly inferred to have originated through innovations, although some mLECA-OGs have non-eukaryotic origins, specifically, there is an important contribution transferred through viruses (15%, 10% from Nucleocytoviricota) among the apoptosis genes (see **Supplementary Table 7**).

*Cellular organisation*

The morphological complexity of unicellular eukaryotes, the mechanical integrity of the cell, the intracellular traffic of molecules with motor proteins and its movement and division depend on the cytoskeleton[55]. The origins of this crucial component of the eukaryotic cell has been deeply studied[56–59], and all the reconstructions, including ours, agree on a complex cytoskeleton in LECA. We detected some of the actins and actin-related proteins (ACTB_G1 - K05692, ACTF - K10355, ACTR1 - K16575, ACTR6 - K11662 and ACTR10 - K16576), all the proteins of the Arp2/3 complex, which is a protein complex with a crucial role in the organization of the cell's cytoskeleton. Our reconstruction of LECA also contains all Tubulins (TUBA to TUBE, K07374, K07375, K10389, K10390 and K10391) and many of the Kinesins. The motor complexes were also present in LECA with high completeness, suggesting an active cargo-based transport. We detected the presence of the dynein, the dynactin and the kinesin complexes. There were some proteins of the myosin complex, albeit this was less complete. These features, as well as the cytoskeleton and the flagellum, are mainly inferred to be innovations (42% - 45% of the proteins), however, some other components were imported from non-eukaryotic organisms (we detected 9% of the cytoskeleton coming from Asgard archaea) (see **Supplementary Table 7**). These results provide insights into the genetic basis of the currently discussed excavate cellular morphology of LECA[4]. The presence in LECA of these genes supporting a well-developed cytoskeleton and cellular structure, would also explain the scattered distribution of the excavate morphology[3,4], and the diverse distribution of eukaryotic morphologies across the eukaryotic phylogeny[60].

*Mitochondria*

The mitochondrion is one of the main defining features of eukaryotes. Earlier studies have defined the metabolic features of the ancestral proto-mitochondrion[61], with the organelle's proteome experiencing deep transformations as the different eukaryotic groups diverged from each other[62–64]. Our LECA reconstruction is consistent with earlier reconstructions[61,62,65] and with the presence of eukaryotic mitochondrial pathways previously established as derived from the proto-mitochondrial ancestor such as the respiratory complexes and most of the subunits of the ATP-sythase, all mitochondrial Fe-S cluster assembly proteins, and most of the components of the mitochondrial versions of ribosomal proteins and DNA and RNA polymerases. Missing components of the above-mentioned pathways are lost due to the stringent phylogenetic support thresholds required in our reconstruction. Similarly, we cannot uncover the presence in LECA of the TatA and TatC translocases and the heme exporters, which have been previously inferred to be present in LECA based on their phylogenetic distribution[66]. Expectedly, due to the required phylogenetic breadth, our reconstruction misses some ancestral mitochondrial pathways that are not widespread across eukaryotic supergroups due to differential loss, such as the SecY and rpoD translocases[67].

*Overall discussion*

The reconstruction of the features of an organism that lived around ~1.8 – 2.4 Bya[68,69] depends on the data, the assumptions and criteria used to define that organism, and the methodology used. We tackled this problem from a broad perspective, trying to infer the basic features of LECA. Our results depict a rather complex organism that resembles more current extant unicellular free-living eukaryotes than its prokaryotic ancestors. However, these reconstructions do not allow us to envisage the actual status of these mechanisms in the past, just their extant features in their descendants. The current versions of the cellular mechanisms result from changes that can explain the incompleteness of some of the reconstructed features. The presence of typical proteins of a prokaryotic process mixed with those found in the eukaryotic one can be the product of a mixture of acquisitions and innovations during the symbiotic origin of eukaryotes. This suggests that, at some point during eukaryogenesis, prokaryotic and eukaryotic steps (genes) of different mechanisms and pathways coexisted and complemented each other. Later, by gene replacement, acquisitions, *de novo* origin of genes and selection, the current eukaryotic mechanisms that we can detect in extant organisms were, eventually, selected.

##### *Comparison with free-living, unicellular, extant eukaryotes*

We benchmarked our reconstruction of LECA against a set of free-living unicellular eukaryotes of different trophic habits: phagotrophs (FLUPs), osmotrophs (FLUOs) and autotrophs (FLUAs), with particular attention to phagotrophs, as it is inferred to be the ancestral lifestyle. We observe (**Fig. 1b)** an overall good agreement of the proportions of COG categories for the inferred LECA proteome and the individual proteomes of FLUPs, despite high variability within extant organisms. Some categories stand out, however, such as a relatively higher contribution of translation and ribosome (J) and some metabolic categories in LECA than in extant eukaryotes, and a relatively higher proportion of signal transduction (T), and cytoskeleton (Z) in extant eukaryotes than in LECA.

These differences are likely due to post-LECA expansion/reduction of the protein repertoire for these functions and probably reflect lineage-specific adaptations. This is especially true for signal transduction, and, to a lesser extent, cytoskeleton and cell cycle, which bodes well with our depiction of LECA as a rather complex organism, but lacking finer regulatory and signaling machinery that is present on extant eukaryotes. Reduction in metabolism categories can be due to the expansion of these categories altering the relative expansion of more housekeeping functions, whose genes could remain stable, or even decrease in number due to lineage-specific specializations. Our aim with this figure is to assess our LECA reconstruction against the backdrop of extant unicellular organisms, and characterizing the innovations and expansions that drove the increase in certain COG categories is beyond the scope of this paper.

##### *LECA acquisition waves*

To detect whether the genes from a specific donor were acquired at the same relative time (a transfer wave), we computed the normalised stem length previously proposed as a measure of relative time for the eight main contributors to the LECA proteome (**Extended Data Fig. 2**). This measure normalises the branch length connecting LECA to the ancestor with its closest non-eukaryotic relative, with no detectable dependence on the functional category of the genes (**Supplementary Fig. 10**), nor the classification as main or additional contributor (**Supplementary Fig. 9**). It normalises it by the median LECA-to-tip distance, obtaining a relative measure of evolutionary time[22,70]. The posterior distributions of the mode, the point of the distribution that we assume as the relative acquisition age, depict at least two main blocks, an archaeal set, which are inferred to have happened before the second set of contributions from bacteria and viruses mainly. The acquisitions from Alphaproteobacteria are always located among the shortest stem lengths, indicating that these transfers happened in a rather late stage of eukaryogenesis after other major bacterial contributions. This conclusion is even stronger when the criterion is more stringent.

To dissect the probability of two waves happening at the same time, we compared the posterior distributions of each pair of donor waves. This resulted in a heatmap of posterior probabilities of the difference between the relative times (**Extended Data Fig. 3**), the higher the probability number, the less likely that both events happened simultaneously. Contiguous waves are difficult to distinguish, as their probability of occurring at different times is low except in some cases (**Extended Data Fig. 3**, see the first upper diagonal of the matrices). This pattern is accentuated among the bacterial donors (with some exceptions), suggesting a continuous bacterial contribution which did not happen in specific temporally spaced waves. The stringent criterion provides higher probability values, as previously commented, the posterior distributions are more spaced (**Extended Data Fig. 2**), and this is translated to a higher probability of them happening at different relative ages.

Our results agree with a symbiotic origin of the eukaryotic cell from an archaeal ancestor, as archaeal genes are inferred to be the oldest ones in the LECA proteomes. Moreover, our reconstructions and relative timing agree with a mito-intermediate or mito-late scenario for the acquisition of the mitochondria as we observe a tendency for shorter stem lengths for the genes with alphaproteobacterial ancestry as previously observed[22,71]. Altogether, we can envisage a eukaryogenesis process with more than one specific bacterial partner. These partners transferred genes at different times, although they may have overlapped, suggesting a strong ecological interaction of the proto-eukaryotic population with other diverse bacterial populations. Although these waves can be used to detect interaction (at some point, both organisms coexisted), they do not allow us to predict which kind of ecological or symbiotic interaction they had. What we can sketch from our data is that eukaryogenesis happened in bacterial-rich environments and in a physical disposition that allowed the interaction. The co-occurrence of bacterial and viral populations with the proto-eukaryotic ones and the acquisition of relevant genes by the latter, allowed the proto-eukaryotes to evolve their cellular complexity.

##### ***Nucleocytoviricota-mediated transfers to LECA***

Our results suggest an important role of viruses during eukaryogenesis, mediating the transfers between prokaryotes and LECA. In our analysis they appeared in numerous trees either as part of the LECA group (513 out of 6,469 directly acquired mLECA-OGs in TOLDBA - 8%), indicating recent bouts of eukaryote to virus transfers, or as present in the sister to LECA, either exclusively or as part of a larger group of non-eukaryotic sequences (1,087 out of 8,790 –12%). This last set of trees are of particular interest given that it means viruses served as intermediaries between the prokaryotic donor and LECA. We can then divide these trees in which ones have viruses as the main donor (the number of viral sequences exceeds the number of other potential donors) or a minor donor (the number of viral sequences is surpassed by another donor). We searched which prokaryotic taxonomic groups appeared together with viruses in a mixed sister group to LECA when they appeared as either minor or main donors. Interestingly, when viruses are the main donors, Myxococcota often appear grouped with them in 48 out of the 558 cases (9%), the second most abundant group is Gammaproteobacteria followed by Margulisbacteria with 35 and 20 cases respectively. The fact that viruses and these bacterial groups appear together in the same clade could indicate that they are involved in recent HGT events. We then checked which viruses were most often found as sister to LECA. Nucleocytoviricota phylum and the Megaviricetes class, a class of nucleocytoplasmic large DNA viruses, was most often present in the sisters, in 78% of the cases (436 out of 558 mLECA-OGs). The most represented family is the Mimiviridae family, which are giant viruses that infect protists (345 out of 558 -

62%). This pattern mimics the one found in viruses grouping within LECA, which also presents a majority of Mimiviridae, though the proportion is slightly lower, 48%.

Among other sources of phylogenetic error, such as long branch attraction, there is the possibility that the grouping of LECA + viruses is an artefact caused by misplacing the root. The algorithm we use places the root at the non-eukaryotic clade with the largest distance to the LECA group. If there are no other prokaryotes that can serve as outgroup, this may cause an artifactual root placement that divides LECA and viruses and causes us to call a relation that otherwise would be interpreted as an HGT event between eukaryotes and viruses. These trees have been classified as of “unknown origin”, joined with those that have a low taxonomic distribution in prokaryotes, due to the complexity of rooting them properly. In these trees we find viruses in 404 out of the 1,763 unknown trees –23%.

Of particular interest are those trees that present a majority of viruses in the sister closest to LECA and have additional prokaryotic sequences grouping to the clade of LECA + viruses. We classified this class of trees as “virus-mediated acquisitions” and are evidence of viruses playing a role in mediating the transference of genes between prokaryotic species and LECA. When analysing the prokaryotic sisters to the LECA + viruses group, the presence of Asgard Archaea stands out (10%, see **Supplementary Table 8**).

We looked into the functionality of the genes presented in these trees, both in terms of assigned consensus KOs and consensus COG functional terms. Regarding the first, 639 trees were assigned a KO. Many of them mapped to similar KOs resulting in 448 unique KOs. The two KO terms that appear most often are protein phosphatase 1L (K17506) and serine/threonine-protein kinase (K08857) that appear seven times each. When relating to COG functional categories the most abundant terms present are related to T: signal transduction mechanisms, O: Post Translational modification, protein turnover and chaperones and J: Translation, ribosomal structure and biogenesis.

***Assessing our results in the context of different models for the origin of eukaryotes***

Our reconstruction of the ancestries (**Fig. 2a-c**) of the genes present in LECA shows a mixture of contributions with blurry functional boundaries (**Fig. 2d,e, Supplementary Fig. 8 and Supplementary Tables 4 and 7**). These contributions stem from a diversity of bacteria that transferred genes to the proto-eukaryote during the whole eukaryogenesis process. In a background of multiple transfers from minor contributors, some major donors can be identified that donated waves of genes, with some transferring earlier than others (**Fig. 3a, Extended Data Fig. 2 and 3**). The number of clades, their relative contributions and the relative timing in which the proto-eukaryote acquired the genes suggest that bacteria, archaea and viruses cohabited in an environment that favoured the close interaction and, eventually, the symbiogenic origin of proto-eukaryotes.

The nature of the environment where the origin of eukaryotes took place has been thoroughly discussed by the authors of the different hypotheses. Some hypotheses are more specific, in which eukaryogenesis happened in microbial mats (syntrophy[72,73] and D. Searcy hypothesis[74] or benthic habitats of shallow oceans –E3 hypothesis[75]). Otherwise, other hypotheses just predict the characteristics of the anoxic nature of the environment (hydrogen hypothesis[76,77] and reverse flow[78]).

From the ancestries of the genes inferred to have been present in LECA, we detect a diverse and continued bacterial contribution during eukaryogenesis (enclosed between the first archaeal contribution and the last bacterial –in our case, the ancestor of the mitochondria– contributions). This is in line with the origin of eukaryotes in a bacterial-rich environment (**Fig. 3b**) and favouring their close interaction, as they may have metabolically complemented during the initial stages of eukaryogenesis and eventually integrated into a unique cell, or took part in transient symbiotic interactions that left a footprint in the form of HGT. Apart from the canonical contributors to eukaryotes (Asgard archaea and Alphaproteobacteria), we detect contributions from Myxococcota and Planctomycetota. Moreover, we observed a significant role of viruses during the origin of eukaryotes, which may have aided the gene transfer process (**Fig. 2a,c**). This distribution of significant gene donors supports the idea that eukaryogenesis could have happened in bacterial-rich environments, and as previously proposed[73], microbial laminated mats can be a candidate environment for this event from our data.

A microbial mat is distributed in three main layers: 1) the oxic and phototrophic zone, inhabited by phototrophs and aerobic heterotrophs; 2) the anoxic and phototroph layer; and 3) the anoxygenic layer, in which the sulfate reduction and methanogenesis happen[79,80]. The biochemical properties of the environment select the bacteria in each zone. Therefore, although the composition is complex and diverse in all the layers, we find different proportions of the bacterial clades between the layers[81–83]. The shallower layer is predominantly cyanobacterial, followed by phototrophic Alpha- and Gammaproteobacteria species, which fix $CO_2$ and synthesise organic matter for the rest of the mat. Some studies also found Bacteroidetes in this upper layer. Underneath, the mats contain anoxygenic phototroph species, such as those of the Chloroflexota and Chlorobi (green sulfur bacteria) clades. Finally, the deepest zone of the mat has sulfate-reducing bacteria (SRB, mainly Myxococcota[84], and both cultured isolates of Asgard archaea are known to participate in sulfur-based syntrophies[75,85]) and methanogens (mainly linked to Thermoplasmatales[83], although methanogenesis is thought to have been ancestral in Archaea and then lost in the archaeal ancestor of eukaryotes). Planctomycetota appears throughout the microbial mat, although its proportion is higher in the anoxic and anoxygenic zones. Moreover, viruses are found in the mats, as well as the CRISPR-Cas systems in the mat prokaryotes[86].

Our relative dating analyses for the main contributors to the LECA proteome (**see Supplementary Discussion, Fig. 3a and Extended Data Fig. 2 and 3**) show a bacterial transfer continuum. However, we can distinguish between early and late transfers. We observe that Asgard archaea and Planctomycetota were early contributors, while we detect that Alphaproteobacteria transferred in a late stage of the eukaryogenesis. Hence, our data lend support and suggest potential partners to the pre-mitochondrial symbiosis hypothesis[87], which posits that, prior to the engulfment of mitochondria, other bacterial symbionts contributed genes to the eukaryotic lineage. As predicted in the syntrophy hypothesis[73], eukaryogenesis may have happened in two stages: first, the symbiosis of a hydrogen-releasing Asgard archaeon and a complex myxobacterial-like bacterium, which would have occurred in the anoxic zone of the microbial mat where they cohabited; and second, the symbiosis with the ancestor of the mitochondria, that could have happened in the upper layers of the mat where Alpha- and Gammaproteobacteria are found to be more abundant[73]. This order of events correlates with the relative ages we found with the stem lengths. Our data suggest the involvement of an additional partner, likely before the myxobacterial-like organism, affiliated to Planctomycetota. The organisms found in the deep layers of the microbial mat are inferred to have contributed earlier than those found in the shallower ones, supporting the prediction of the syntrophy hypothesis.

The main outcome of this study is that the ancestral genome of LECA had contributions from diverse donors. Therefore, our data suggests that eukaryogenesis is more likely to have happened in environments with a complex community structure of prokaryotes and also viruses. This strengthens the idea of a syntrophic origin of the eukaryotic cell. According to our reconstructions it is difficult to establish a specific syntrophic scenario, and several proposed ones are compatible with our results as we observed the proposed metabolic potentials in our inferred donors. Other models are harder to reconcile with our observations. In addition, new models that may combine features with existing ones, may better explain the overall observations.

Regarding the nature of the archaeal host, early theories suggesting a methanogenic archaeon seem to be discouraged by our data. Despite an ancient origin of methanogenesis in the archaeal TOL, the last archaeal-eukaryotic common ancestor (LAECA) is inferred to not have been methanogenic[57] based on the inferred absence of key methanogenic enzymes (coenzyme M methyltransferase (MtbA, K14082), and methyl-coenzyme M reductase (MCR, K00399, K00401, and K00402), see “Biomass and energy conservation from formate, trimethylamine, and formaldehyde” on their Supplementary Discussion for the rationale behind the usage of these genes as markers for methanogenic potential). Our results also show that, despite prevalence of some of the modules related with methanogenesis, among the 60 genomes used to infer the Asgard archaeal donor, 37 have MtbA and only 8 have at least one unit of the MCR complex. We searched these genes also in the 12 selected genomes that belong to Heimdallarchaeota, and the results agree with the Asgard archaeal donor ones, none of the genomes showed complete MCR (2 have at least 1 KO) and just 2 of them have MtbA.

Among the proposed metabolic scenarios, the Hydrogen Sulphur-based (HS) syntrophy[73] argues for an initial incorporation of a hydrogen-producing Asgard archaeon (future nucleus) within a sulphate-reducing deltaproteobacterial (Myxococcota) host, with a subsequent acquisition of the facultatively aerobic, sulphide-oxidizing alphaproteobacterial ancestor of mitochondria. Our results provide support for the three partners (and additional ones). Moreover, the key metabolic properties assumed by this model are present. This is the existing metabolic scenario that finds more support with our data, along with the E3 hypothesis. The E3 hypothesis proposes an organic compound degrading Asgard archaea, an SRB, and an aerobic organotrophic mitochondrial ancestor. For this hypothesis we are able to detect the proposed metabolic properties for archaeal, bacterial and mitochondrial ancestors (with varying prevalences, such as 0.63-0.67 for fermentative metabolisms in the archaeal ancestor, 0.44 and 0.5 for sulfate reduction for the Myxococcota and Planctomycetota ancestors that may have cohabited with the protoeukaryote, respectively, and overall high prevalences for the organotroph ancestor of the mitochondria).

Other metabolic scenarios find less support from our data. Although all of them involve an Asgard archaea and an Alphaproteobacterial (or related) ancestor, which we find evidence for (among others), some of the metabolic properties required by these models are at odds with our reconstructions. For instance, the Searcy syntrophy hypothesis[74], which involves a sulfur-respiring archaeon and an $H_2S$-utilising bacterium, contrasts with the absence of sulfur metabolism in our reconstruction of the Asgard archaeal ancestor. The revised hydrogen hypothesis[76,77] proposes an autotrophic Asgard archaeon with the H(4)MTP-dehydrogenase, although we do not find it in our reconstruction of the donor (prevalence 0.02). However, the mitochondrial ancestor proposed in this hypothesis seems likely, as we infer the presence of fermentative metabolism. The reverse flow hypothesis[78] proposes a fermentative archaeal ancestor, for which we detect ethanol and butyrate fermentations with a non-negligible prevalence, however, the Fe-Fe- Ni-Fe- and Fe-only hydrogenases are not detected in the descendants of the mitochondrial ancestor.

Beyond contrasting existing models, our results can stimulate adjustments to them, and even the development of novel scenarios to better accommodate the observations. For instance, given the similarities of the inferred Planctomycetota ancestor metabolism with the inferred properties of the Deltaproteobacterial host of the revised HS syntrophy hypothesis, this hypothesis could be revised by changing the host to a Planctomycetota, to accommodate our relative timing results. This would bring some attractive features of Planctomycetota, such as the complex membrane invaginations[88], and their ability to engulf other cells[88,89]. However, other scenarios, considering the inferred partners and metabolic properties may also be equally plausible. Further research is needed to elucidate eukaryogenesis and there are still many open questions such as the biochemical feasibility of the syntrophic models from an ecological perspective, the plausibility of a membrane shift, and the co-occurrence of the current descendants of the donors in different ecological niches, and the characterisation of their symbiotic interactions.

The potential roles of viruses during eukaryogenesis have been extensively discussed earlier[90–93]. Proposed viral contributions to eukaryogenesis range from a potential source of key genes[94,95], to a direct role in the origin of the cell nucleus either via evolution from viral cell factories[90,96], or as a shelter to protect against viral infections[97,98]. Our results support a major role of viruses in the conformation of the LECA proteome, and point to nucleocytoviridicota –a viral clade often considered in the above discussions– as the key viral clade. Our data indicate that the LECA proteome included a sizable fraction of protein families whose origins can be traced back to viruses. For some of these we can trace farther ancestries from diverse phyla, including Asgard archaea and the major bacterial donors identified in this work, indicating that viruses mediated the acquisition of genes from other ultimate donors. This mechanism could explain part of the breadth of taxonomic origins identified in this work and earlier studies[22]. Additionally it opens the possibility that many of these viral-transferred genes originated from coexisting but now extinct pre-LECA ancestral eukaryotic lineages, which may have facilitated the recruitment of eukaryotic innovations or prokaryotic acquisitions from these proto-eukaryotic donors. Although many of the viral transferred genes have nuclear functions, many others do not, and our results cannot specifically support or refute any of the proposed mechanisms for the origin of the nucleus through viral interactions. Nevertheless both ideas are compatible with our observations.

***Comparison with previous studies***

To assess the effect of the database on our results with respect to previous studies, we compared the taxonomic coverage of our study with that of Pittis and Gabaldón (2016)[22] and Vosseberg et al. (2021)[71]. Our database contains 65,703 prokaryotic proteomes and 185 eukaryotic proteomes. Pittis and Gabaldón's dataset comprises 1,695 prokaryotic and 37 eukaryotic proteomes, and Vosseberg et al. one accounts for 3,457 prokaryotic and 209 eukaryotic proteomes. Moreover, we included 172,004 viral proteins, which were not included in any of the previously mentioned datasets. Regarding the taxonomic coverage of eukaryotes, we retrieved several proteomes for groups that were not considered previously, allowing us to better characterise the LECA node. Whereas Pittis and Gabaldón (2016) and Vosseberg et al. (2021) eukaryotic datasets were focused on Opisthokonta (43% and 58% of each dataset, respectively), our most represented group is the TRASH group (46% of our dataset). The main difference lies in the fact that TRASH is a very diverse group containing several eukaryotic clades (telonemids, rhizarians, alveolates, stramenopiles and haptophytes), justifying its strong presence in the database. Thus, our study covers most of the known diversity in all the domains of life, it includes viruses, and it provides an unbiased towards Opisthokonta framework for LECA.

Regarding possible taxonomic re-classification driving differences in the assignment of contributors, we compared the 692 taxonomic levels employed in Pittis and Gabaldón (2016) with the GTDB taxonomy. We further assessed the affiliation of their genus at phylum level in the taxonomic framework employed by Pittis and Gabaldón 2016 (based in NCBI taxonomy) and the taxonomic framework of GTDB. Most of the phyla have a 1:1 correspondence, with only some exceptions (GTDB phylum "Thermoproteota" encompassing TACK, or Nitrospirae being split into different phyla within GTDB framework) and only a small minority of genomes (taxa) changing phyla. Within phylum Proteobacteria, most alphaproteobacterial sequences remain Alphaproteobacteria within GTDB, beta- and Gammaproteobacteria are joined into class Gammaproteobacteria, Zetaproteobacteria remains the same and the delta/epsilon subdivisions are split into different classes and phyla according to the GTDB species phylogeny. For instance, Myxococcota was previously placed within Deltaproteobacteria, while currently is considered a phylum. Thus, we consider that differences in taxonomic criteria are unlikely to drive major differences in the results, as they do not dramatically affect the discussed groups. As taxonomic modifications are adopted to better reflect evolutionary relationships, our analysis should be considered more accurate regarding taxonomic assignment as compared to that of Pittis and Gabaldón (2016).

We further assessed the difference between our algorithm for donor detection the one used in Pittis and Gabaldón (2016) by re-analyzing the trees of that study with the algorithm used in this one. We calculated the proportion of trees pointing out to each donor by identifying the donor using the MRCA (as in Pittis and Gabaldón) or the most abundant taxon in the sister (as used in this study) (**Supplementary Fig. 7**). The main difference lies in the taxonomic resolution of the assigned clade. While the method used in this study can identify a larger proportion of specific clades, the MRCA approach used in Pittis and Gabaldón assigns more trees to broader taxonomic levels, such as, Bacteria or Archaea (**Supplementary Fig. 7a**). The most abundant taxon in the sister approach accounts for the uncertainty linked to a given taxonomic association. We retrieve the proportion of each taxon, and assign the most abundant, in this case, the average proportion is 0.75, and the median proportion 1. Meaning that the assignment of a specific donor by MRCA may be affected by a few sequences that are the result of pre-LECA intra-prokaryotic transfers. This method allows us to account for the uncertainty associated with an assignment, and recover more specific donors.

We also compared the number of inferred innovations to those predicted in other studies. Vosseberg et al (2021) estimate 2260 out of the total 10,233 gene families to be innovations (22%), which is much less than our estimate of 33% of innovations on average (e.g., 4,219 out of 13,009 in TODLBA). This could be attributed to our filtering approach that may disregard protein similarities restricted to a very minor region (i.e., small domain) of the protein sequence. On the other hand, Hartman and Fedorov in ref.[99] estimated 347 gene families as Eukaryotic Signature Proteins (ESPs), whereas we detect 918. We attribute the increase to an extended sampling of the eukaryotic Tree of Life that allows us to detect proteins of ancient origin that have been overlooked due to their absence in model organisms (so-called jotnarlogs).

### Extended Data

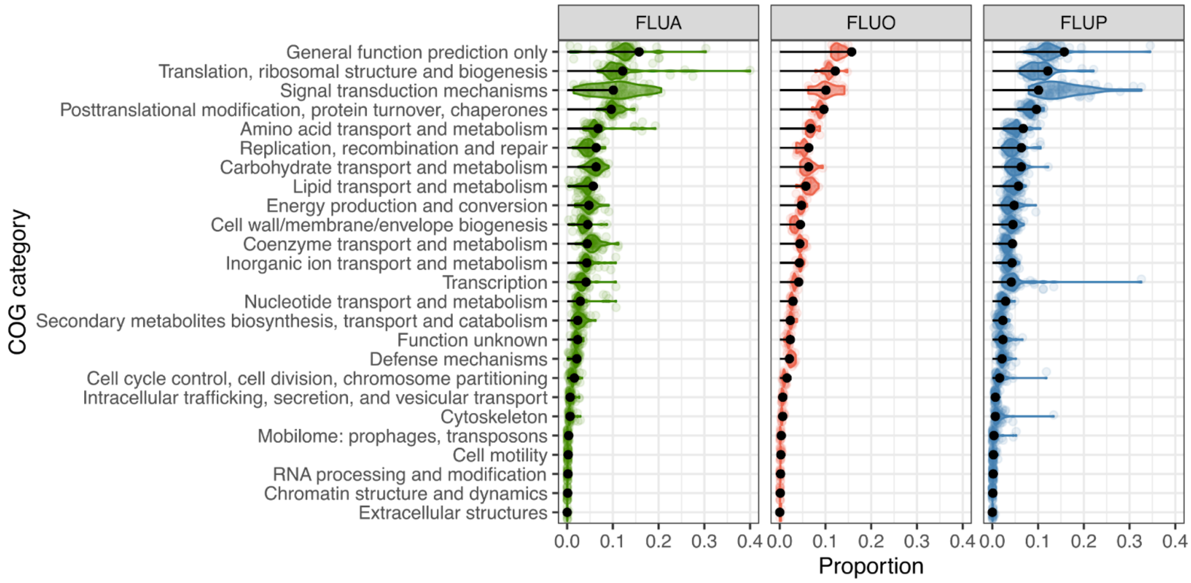

**Extended Data Fig. 1. COG categories distribution for free living unicellular eukaryotic genomes with different trophic strategies**. Each panel shows a trophic strategy, FLUA: autotrophs, FLUO: osmotrophs and FLUP: phagotrophs. Each point refers to a genome with a given trophic strategy and the black lollipop shows the proportion of that category in the consensus proteome of LECA.

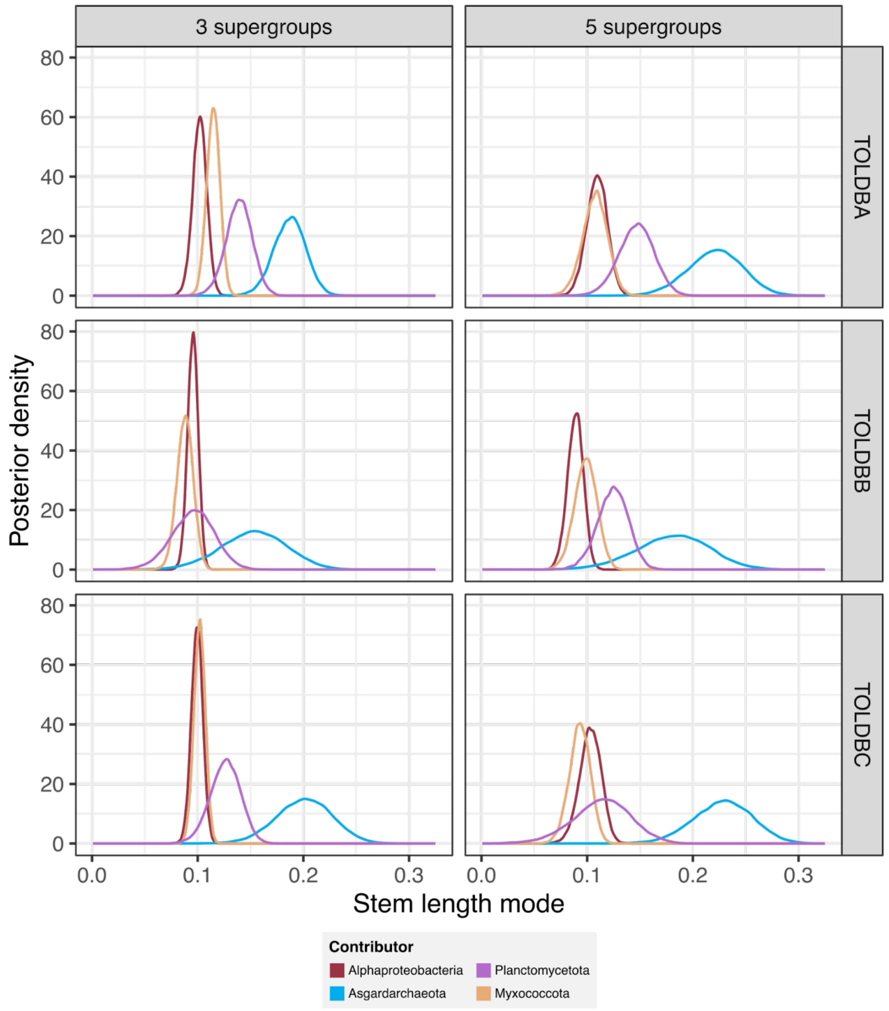

**Extended Data Fig. 2. Acquisition waves in LECA**. Posterior distribution of the mode of the stem length distribution for each taxonomic affiliation. The panels show the results from different databases (in rows) and selection criteria (in columns).

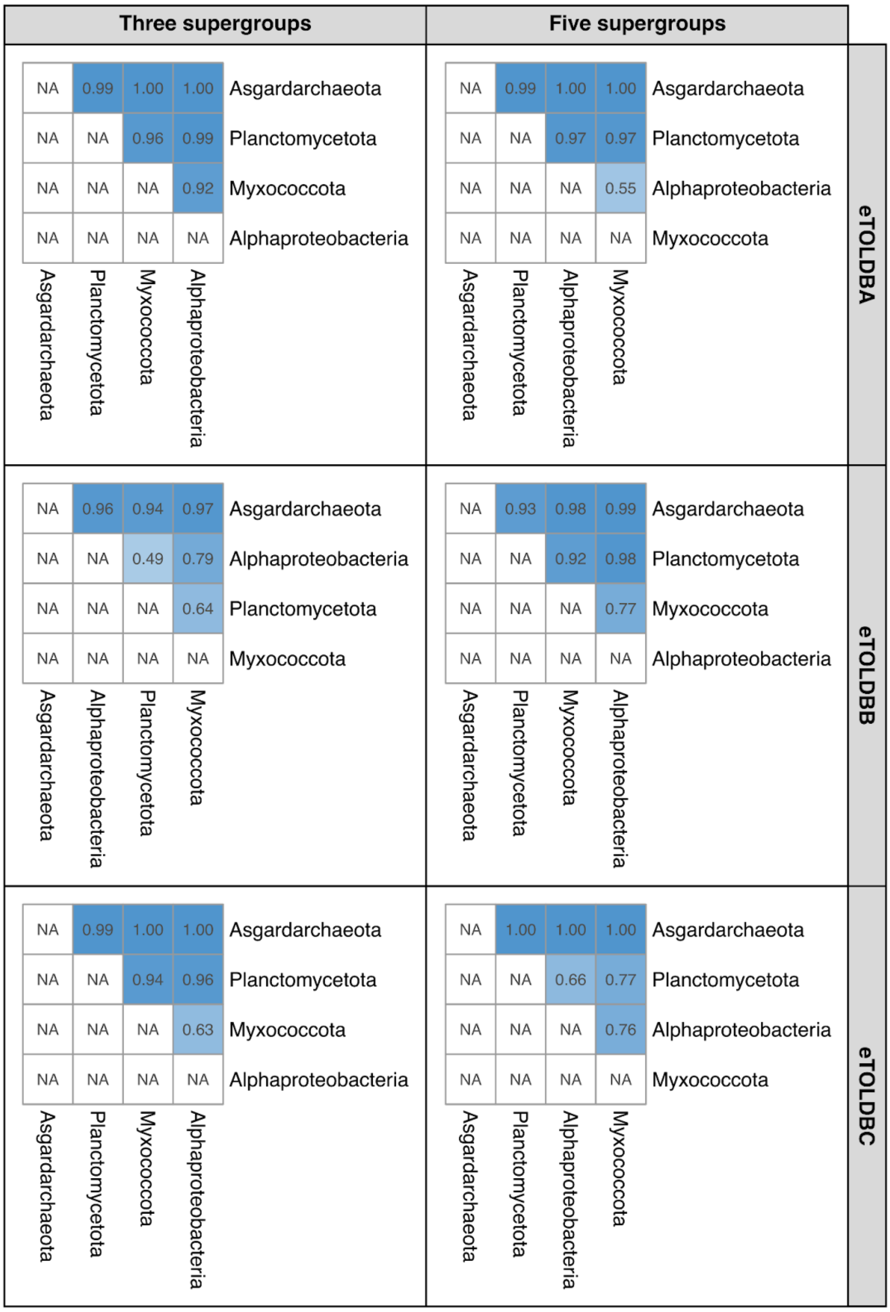

**Extended Data Fig. 3. Probability of the waves occurring at different evolutionary relative time points**. Each heatmap value shows the pairwise probability of the wave from the donor in the row happening before the wave of the donor in the column. The donors are sorted by the acquisition time from older to more recent.

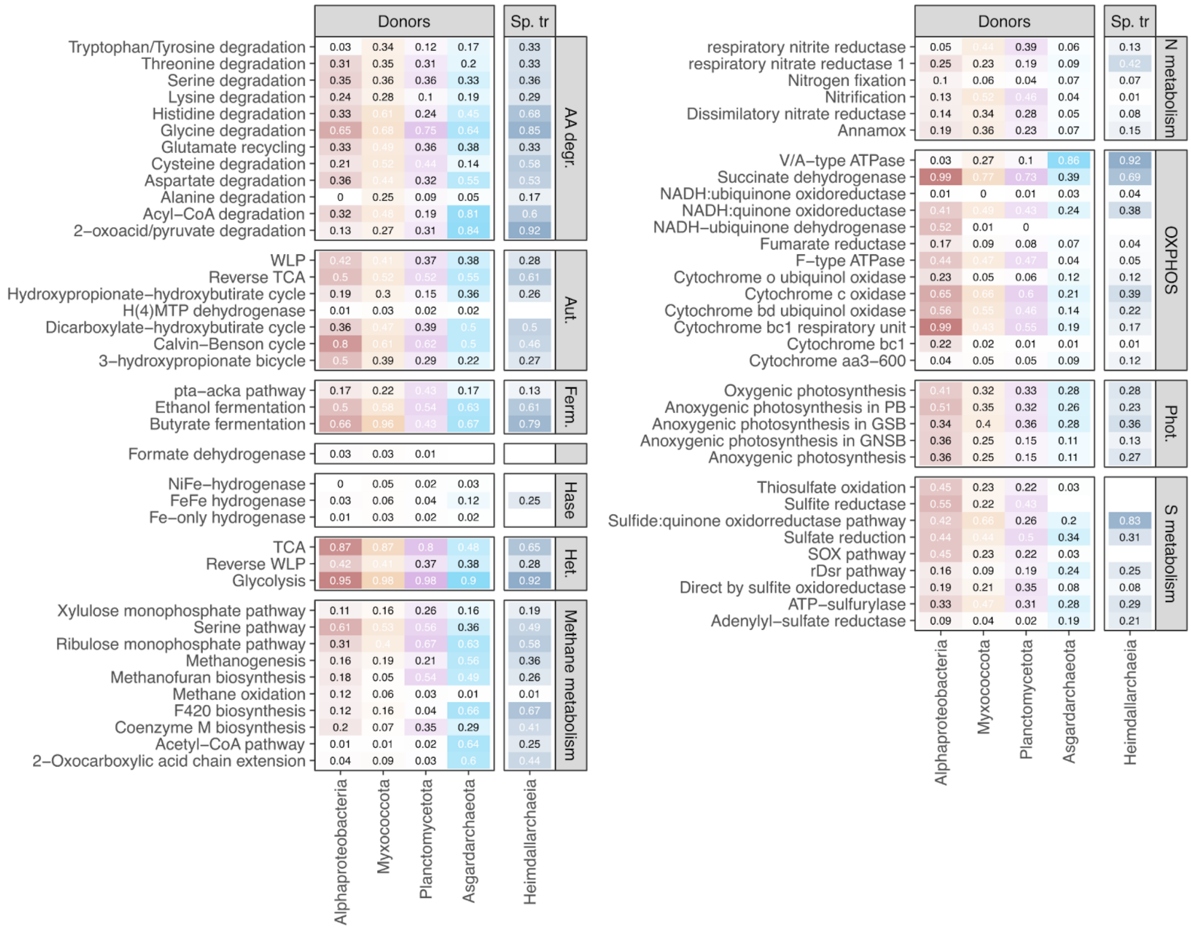

**Extended Data Fig. 4. Prevalence of pathways and enzymes in the genomes of the donor's descendants**. Each box shows a group of metabolic pathways and enzymes. The opacity of the colour shows the prevalence (the percentage of genomes with the feature) of that specific pathway or enzyme in extant genomes from the donor's clades that share at least 50% of the KOs that these donors transferred to LECA. Abbreviations, AA degr.: amino acid degradation; Aut.: autotrophy, Ferm.: fermentation; Hase: hydrogenases; Het.: heterotrophy; N metabolism: nitrogen metabolism; OXPHOS: oxidative phosphorylation; Phot.: photosynthesis; S metabolism: sulphur metabolism.

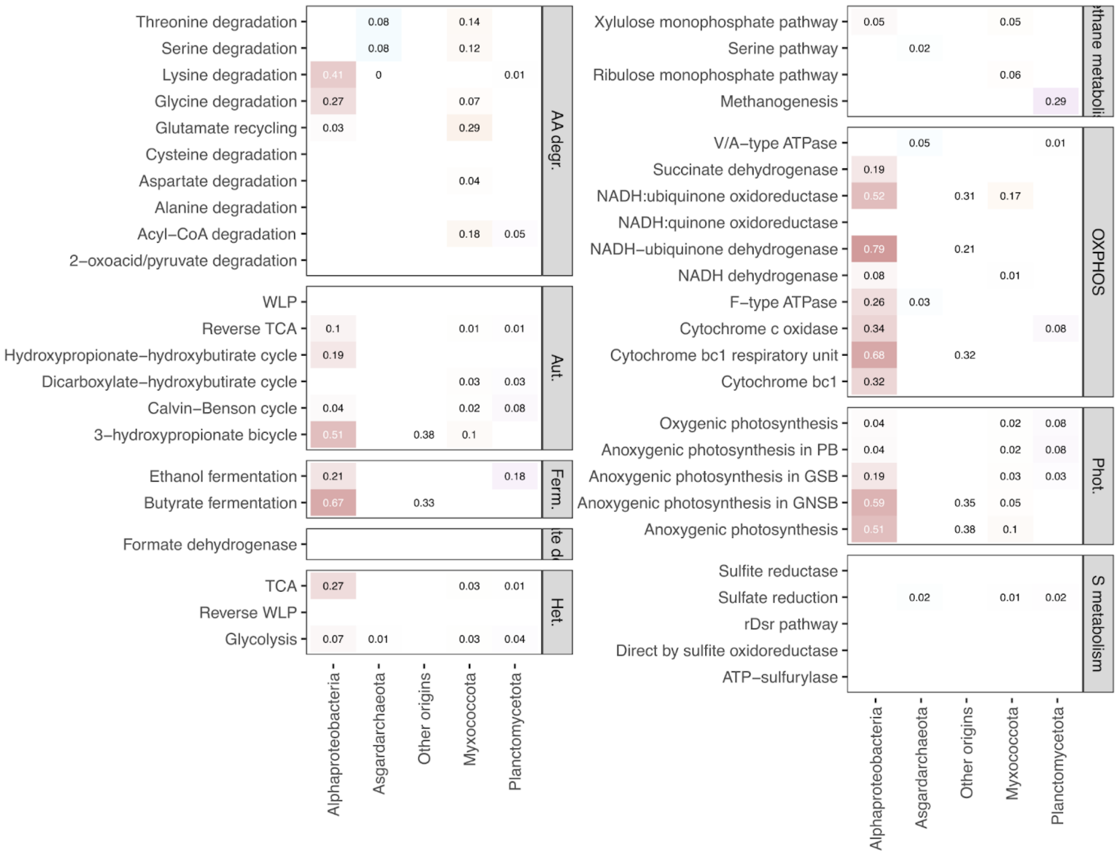

**Extended Data Fig. 5. Origins of the pathways and enzymes in the consensus proteome of LECA**. Each box shows a group of metabolic pathways and enzymes. The opacity of the colour shows the origin's proportion. For each KO, we calculated the proportion of origins for all the databases assigned to a clade, and weighted to the total number of KOs in the module, resulting in a proportion for the feature. Abbreviations, AA degr.: amino acid degradation; Aut.: autotrophy, Ferm.: fermentation; Het.: heterotrophy; OXPHOS: oxidative phosphorylation; Phot.: photosynthesis; S metabolism: sulphur metabolism.

#### Supplementary Figures

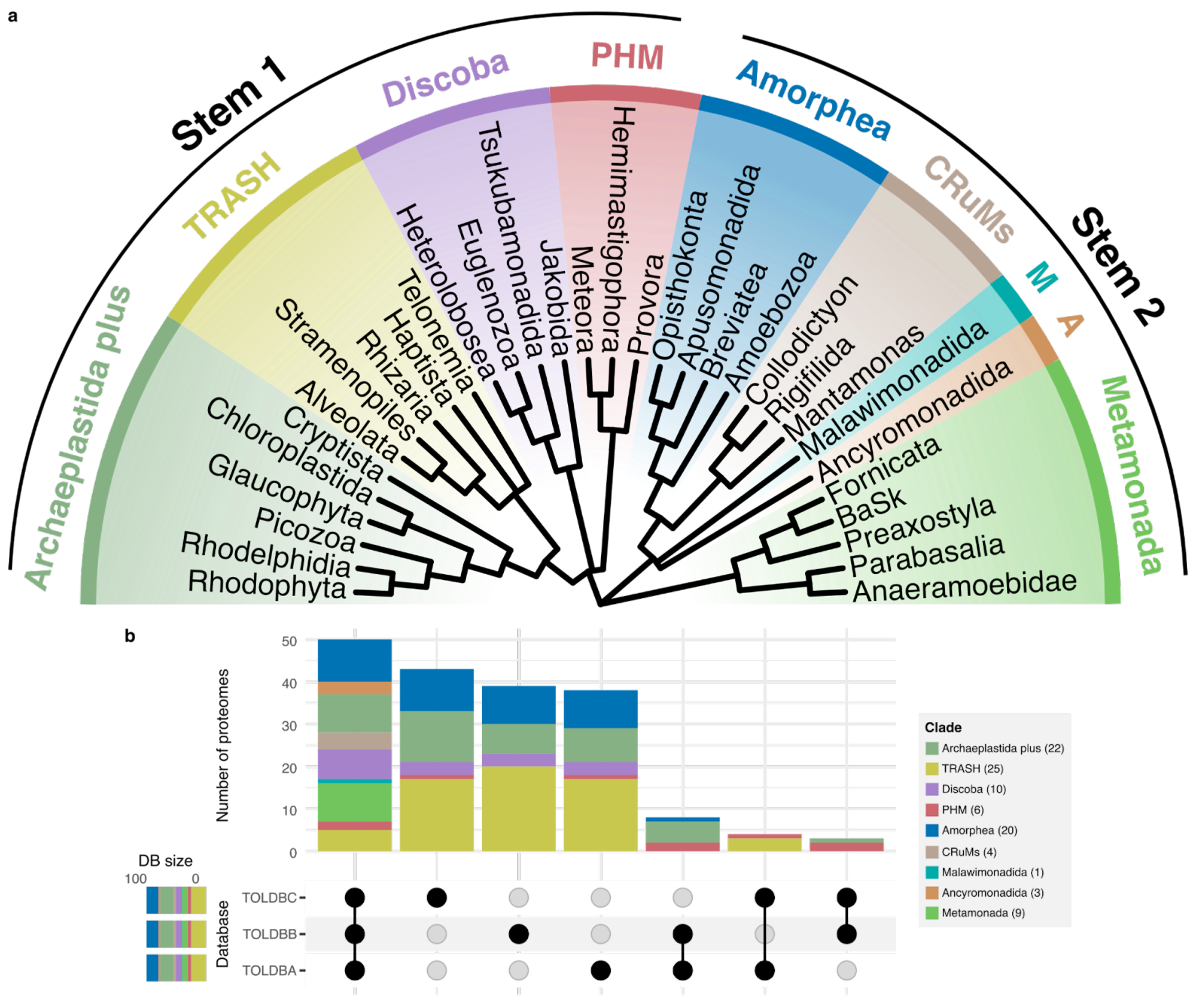

**Supplementary Fig. 1. Eukaryotic backbone tree**. Outer arcs show the two stem groups considered (Diphoda+, and Opimoda + Metamonada). The coloured monophyletic groups indicate the nine considered supergroups. Malawimonadida and Ancyromonadida are considered independent supergroups here as their monophyly is not fully supported. The tips of the tree correspond to the considered divisions. The topology is based on Lax et al., Tikhonenkov et al. and Torruella et al.[1–3], the stems are supported by the recently published root of the eukaryotic phylogeny[4], see supplementary Methods for additional details. b) Upset plot showing the overlap between databases and the distribution of clades in the different dataset intersections. The legend shows the number of proteomes of each eukaryotic supergroup included in each eTOLDB.

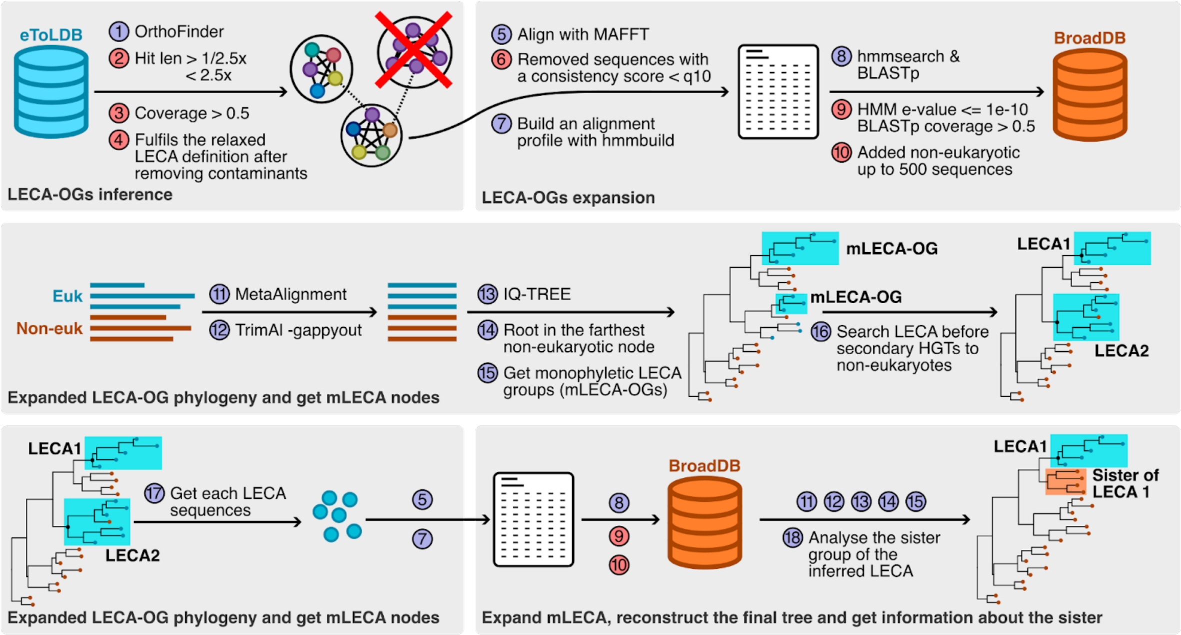

**Supplementary Fig. 2. Graphical summary of the mLECA-OG trees reconstruction**. Each block is in a grey frame with a title at the bottom. Each step of the pipeline is numbered, if the number appears without text, it means that we repeated the specified step with the new data. The blue dots are computational steps, and the red ones are filtering steps. We did this pipeline for each eTOLDB.

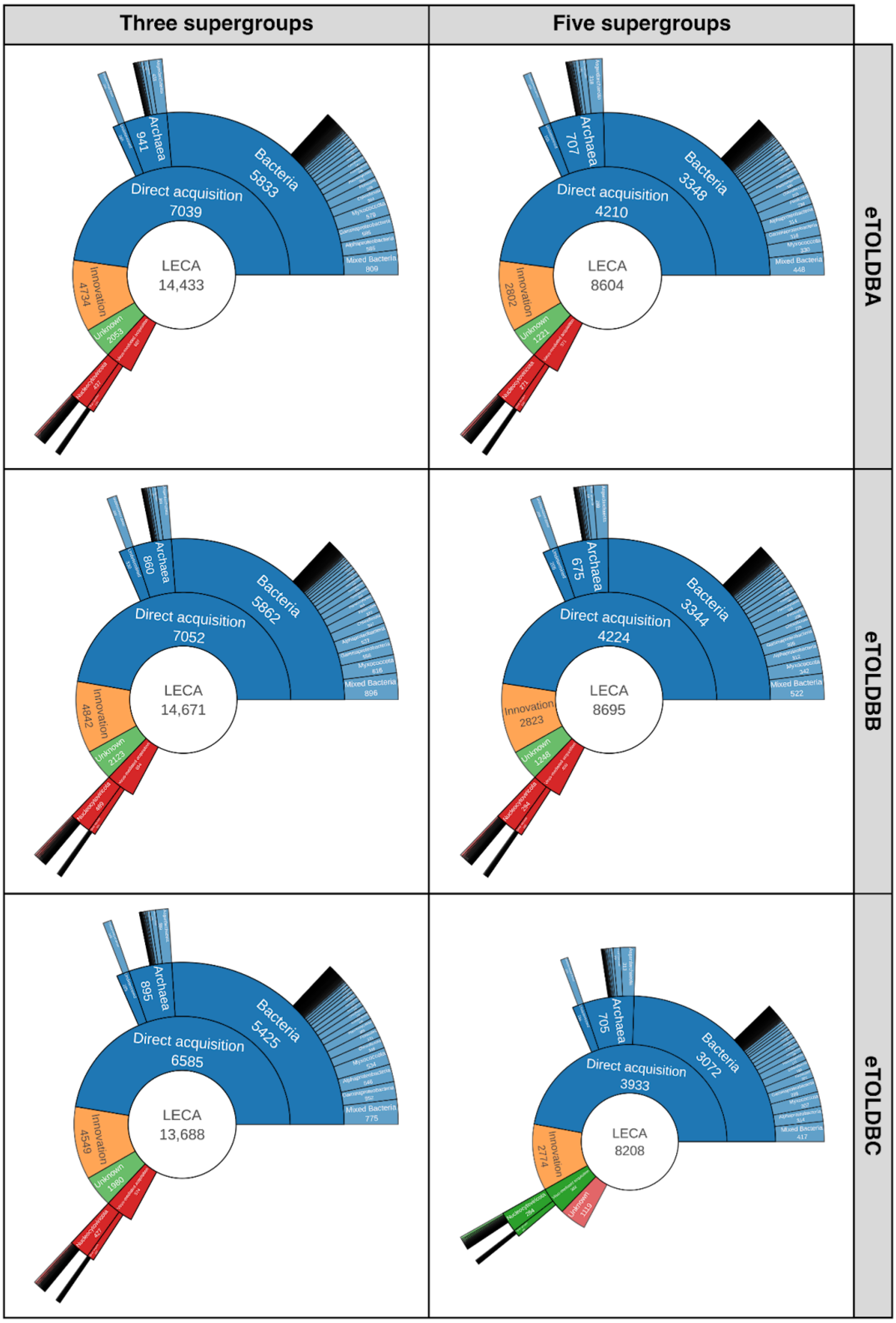

**Supplementary Fig. 3. Sunburst of the origins of the genes inferred in LECA for each dataset**. Each box shows the origins of the mLECA-OGs at the Phylum rank (except for proteobacteria, for which we show the Classes) for each combination of the TOLDB version and the relaxed (3 supergroups) or strict (5 supergroups) criteria for the definition of LECA.

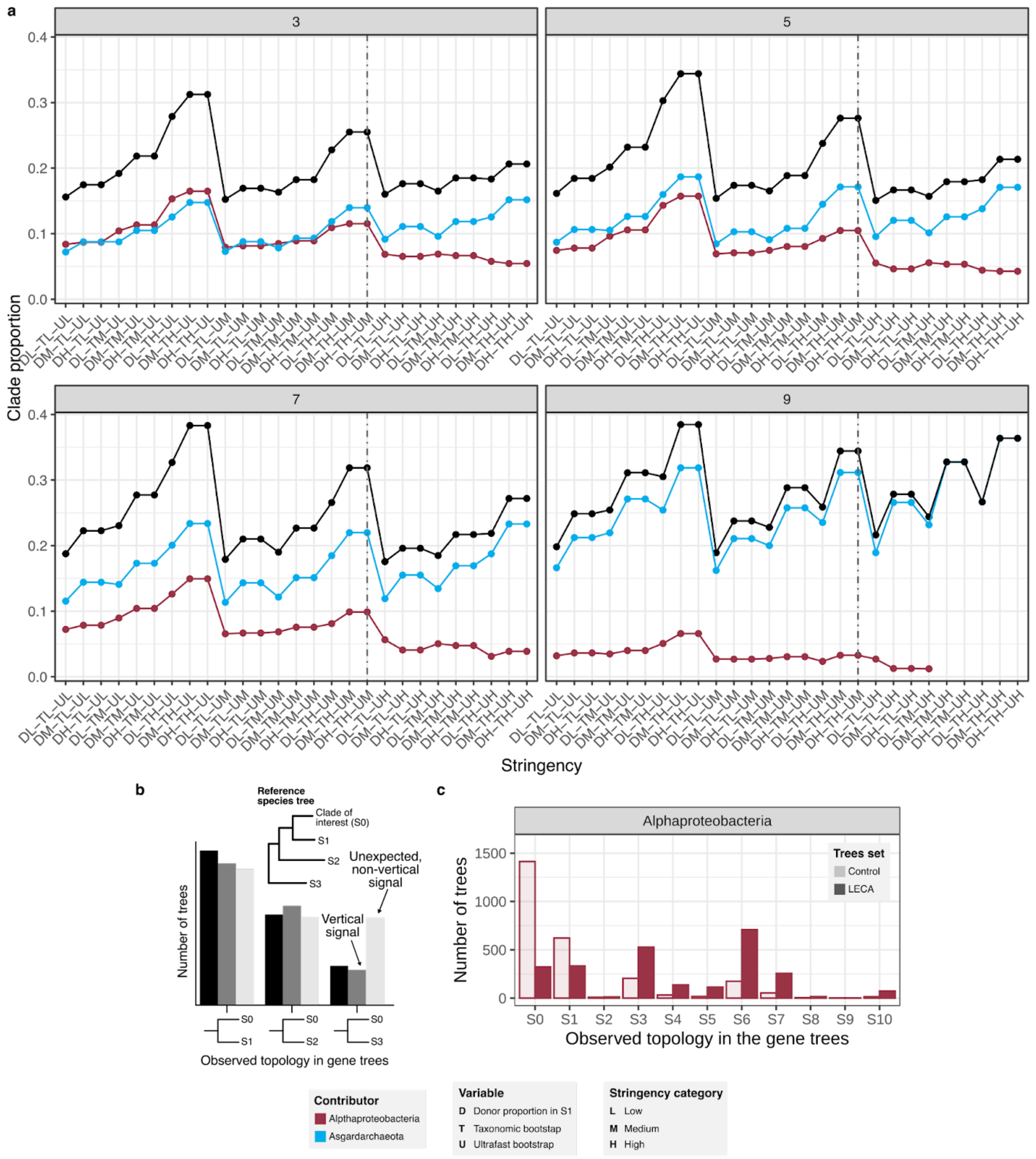

**Supplementary Fig. 4. Impact of stringency thresholds in the proportion of trees with Asgard archaeal and alphaproteobacterial sisters**. a) Stress test threshold for the inference of donors. The blue and red lines show the proportion of Asgard archaea and Alphaproteobacteria in the subset of trees that fulfil the criteria specified in the x axis. The black line shows their sum, the total contribution of Asgard archaea and Alphaproteobacteria to the LECA proteome using a given stringency. The criteria are shown in the legend in the following order: UfBS, taxonomic bootstrap and the donor support. Each panel shows the number of groups required to define the LECA group. The stringency is sorted from lower (left) to higher (right). The vertical dashed line shows the thresholds that we used to detect the most prevalent donors beyond Alphaproteobacteria and Asgardarchaeota. b) Schematic representing the verticality test. On the top is a schematic representation of a species tree with the Clade of interest and the different sisters marked as S1, S2 and S3. The barplots indicate the number of gene trees that have, as sister to the clade of interest, each of the sisters defined in the species tree. Black bars indicate the theoretical signal found in gene trees reconstructed for the main donors, the dark grey bars indicate a set of sisters to LECA that follow the same pattern as the control bars and as such represent vertical inheritance of those sisters. Finally, the light grey bar shows a set of sisters to LECA where the last bar clearly deviates from the control bar and is therefore expected to have evolved through non-vertical evolutionary mechanisms. c) Conceptual scheme and results for the vertical signal found in Alphaproteobacteria families as compared to the LECA donor signal. The lighter bar shows the number of trees that have the indicated species tree sister as sister to the control family. The darker bar shows the number of mLECA-OGs that have clades of that given sister as donors. The decreasing trend and correlation between the lighter and the darker bars would show that the genes evolve vertically. S0 represents the number of trees analysed in Alphaproteobacteria analysis. Then, the different sisters contain the following taxonomic groups: S1 - Gammaproteobacteria; Magnetococcia; Zetaproteobacteria, S2 - Campylobacterota; Aquificota;; Deferribacterota, S3 - Desulfobacterota; Myxococcota; Nitrospirota; Bdellovibrionota; Methylomirabilota; Nitrospinota; SAR324; UBA10199, S4 - Acidobacteriota, S5 - Bacteroidota; Gemmatimonadota; Marinisomatota; Fibrobacterota, S6 - Actinobacteriota; Firmicutes; Patescibacteria; Cyanobacteria; Chloroflexota; Omnitrophota, S7 - Verrucomicrobiota; Planctomycetota, S8 - Chlamydiota, S9 - Dependentiae, S10 - Spirochaetota.

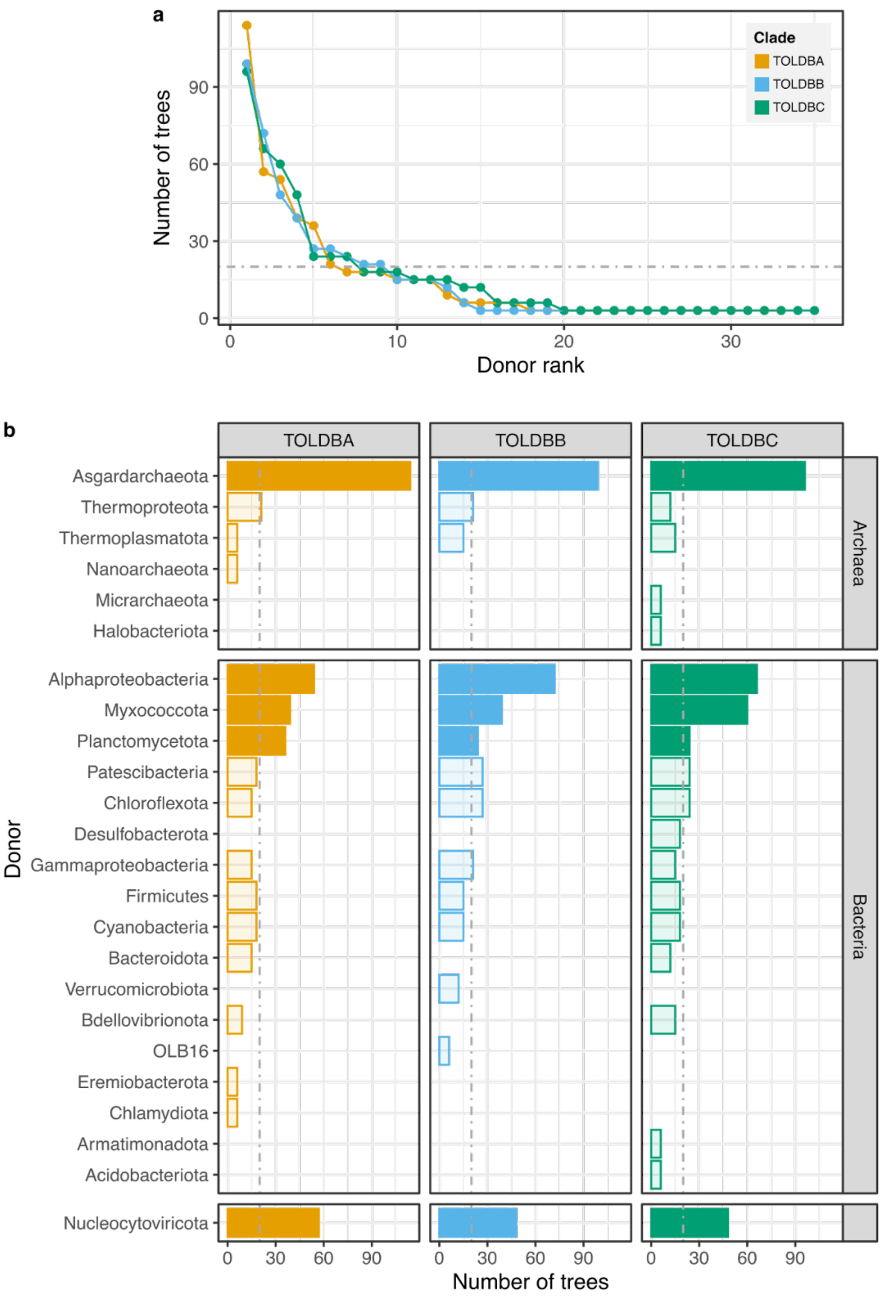

**Supplementary Fig. 5. Non-negligible contributions beyond Alphaproteobacteria and Asgardarchaeota**. a) Number of highly reliable trees (see **Supplementary Discussion**) for each putative donor, with donors sorted along the y-axis by the number of trees and independently for each database. The horizontal dashed line shows the threshold used to identify major donors beyond the established donors Alphaproteobacteria and Asgardarchaeota. b) Number of highly reliable trees assigned to each donor, full bars indicate selected donor groups (which pass the thresholds in all three TODB datasets.

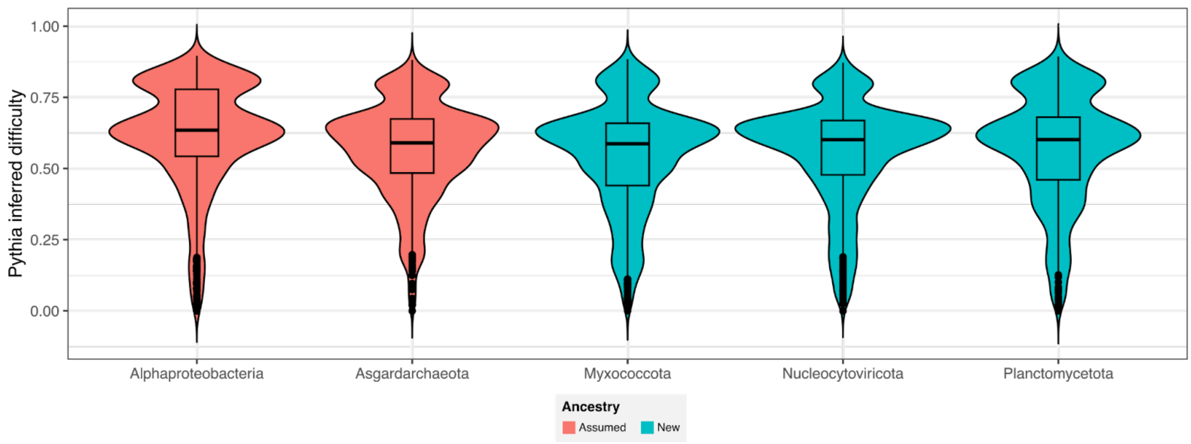

**Supplementary Fig. 6. Distribution of difficulty values of the trimmed alignments per inferred donor as assessed by Pythia**. Violin plots depict the output values of the ML algorithm Pythia, which estimate the difficulty of the tree inference from 0 to 1, where 1 represents a difficult MSA. Statistical significance obtained by pairwise one-tailed (alternative = less) Wilcoxon rank-sum tests against Alphaproteobacteria and Asgardarchaeota, p-values against Alphaproteobacteria 1 (Myxococcota), 1 (Nucleocytoviricota), 1 (Planctomycetota); p-values against Asgardarchaeota 0.983 (Myxococcota), 0.63 (Nucleocytoviricota), 0.099 (Planctomycetota) .

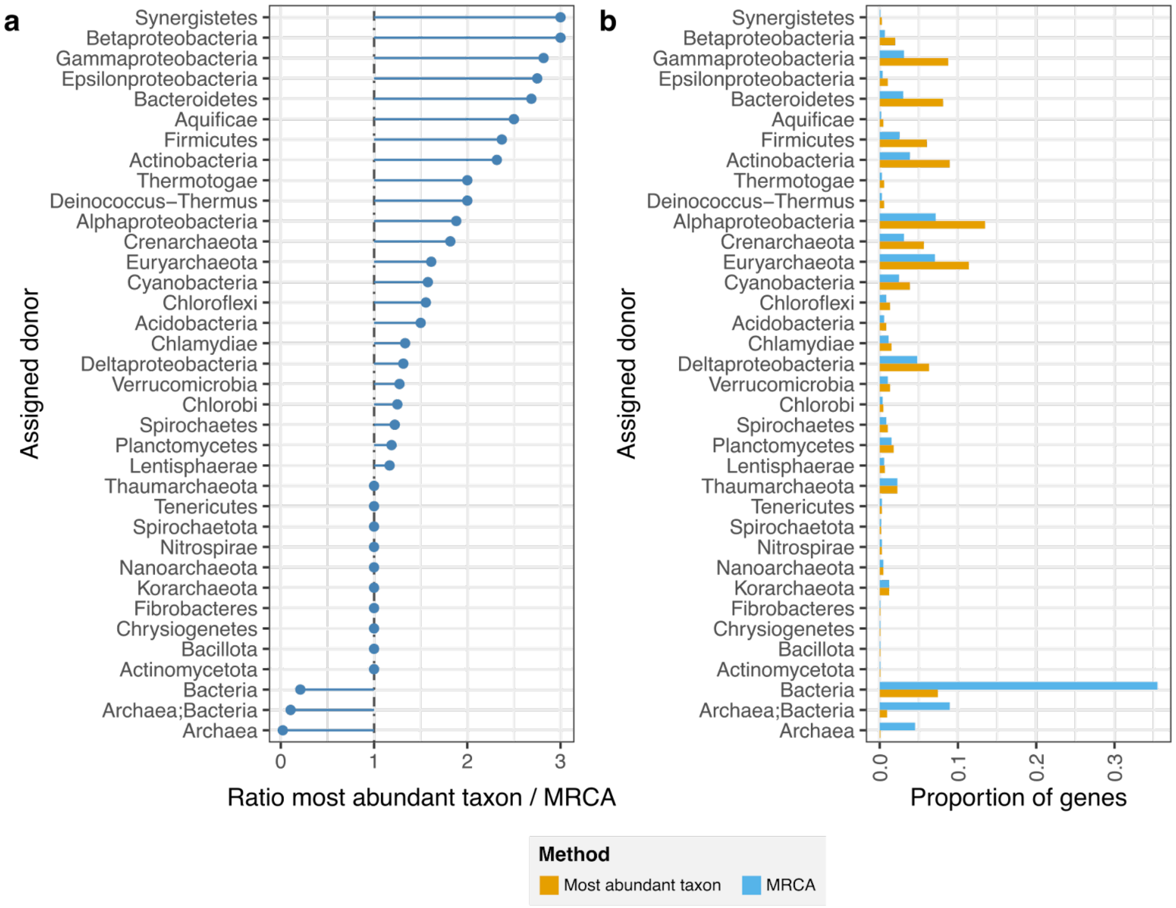

**Supplementary Fig. 7. Comparison of donor assignment methods applied to the Pittis and Gabaldón dataset.** a) The ratio between the proportion of trees assigned to a specific donor using the most abundant taxon method (used in this paper) and the MRCA method (used in Pittis and Gabaldón (2016)[22]). Values higher than one mean that this clade has been assigned more often using our method, and lower than one mean that the clade has been assigned more often using the MRCA. b) Proportion of genes assigned to each donor using both methods.

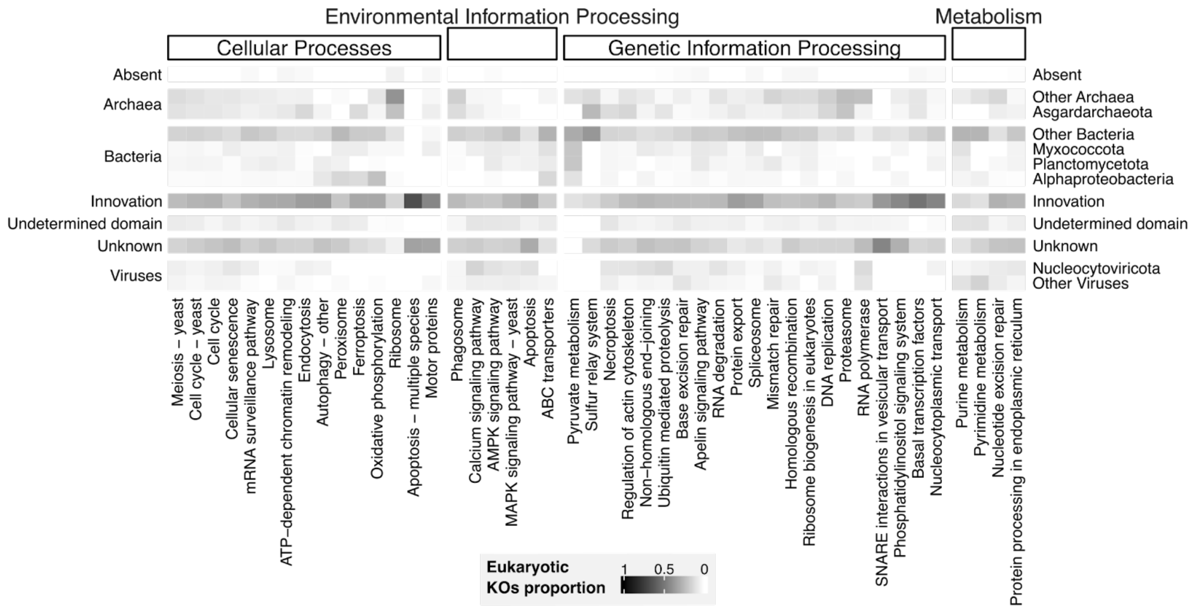

**Supplementary Fig. 8. Relative origin and absence of the main cellular processes features inferred in LECA**. Heatmap representing the proportion of KOs from a certain origin in a given KEGG cellular mechanism. The absent row shows the proportion of KOs we could not infer to be present.

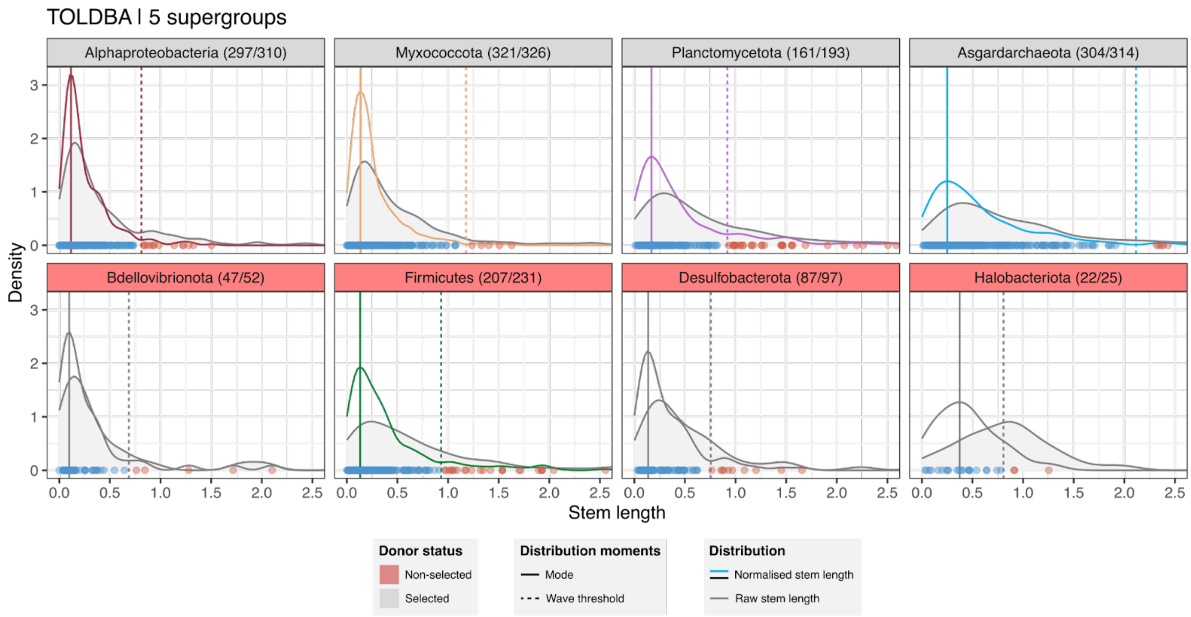

**Supplementary Fig. 9. Raw and normalised stem length distributions for the selected donors and a set of non-selected contributors**. Each panel shows a non-eukaryotic clade inferred to have transferred genes to LECA. The title of the panel shows the clade and the number of genes following the structure: (genes inside the main acquisition wave / all the transferred genes), when coloured in red the clade did not pass our filters. The coloured and black density lines show the normalised stem length distribution, and the grey and filled distributions are those for the raw stem length. Continuous vertical lines show the empirical mode of the normalised stem lengths, and dashed lines show the threshold used to identify the main transfer wave. Dots show the normalised stem length values for individual trees, in blue those inside the wave, in red those excluded from the wave.

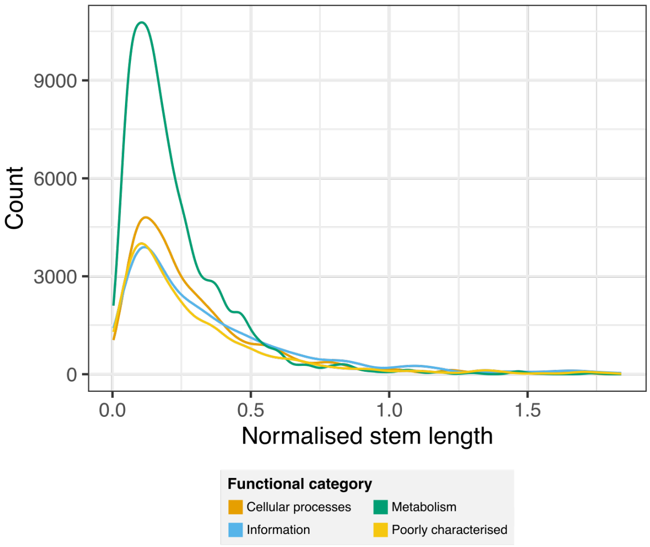

**Supplementary Fig. 10. Normalised stem length distributions per each functional group**. Distribution of the normalised stem length in all the datasets assigned to each function.

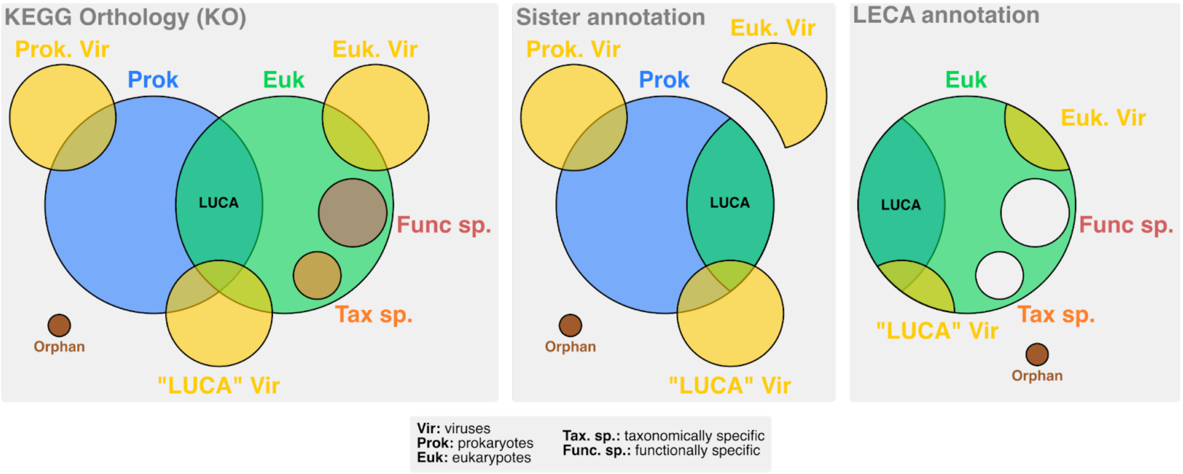

**Supplementary Fig. 11. Functional annotation sets**. Conceptual Venn diagrams of KOs used to filter the protein annotations in the sister and the mLECA group. The KEGG Orthology (KO) database accounts for orthologous groups of different taxonomic and functional deepness. The total set of orthologs is shown on the first panel. The set of KOs used to annotate the sister group is in the middle. The KOs set to annotate the LECA group is on the right panel.

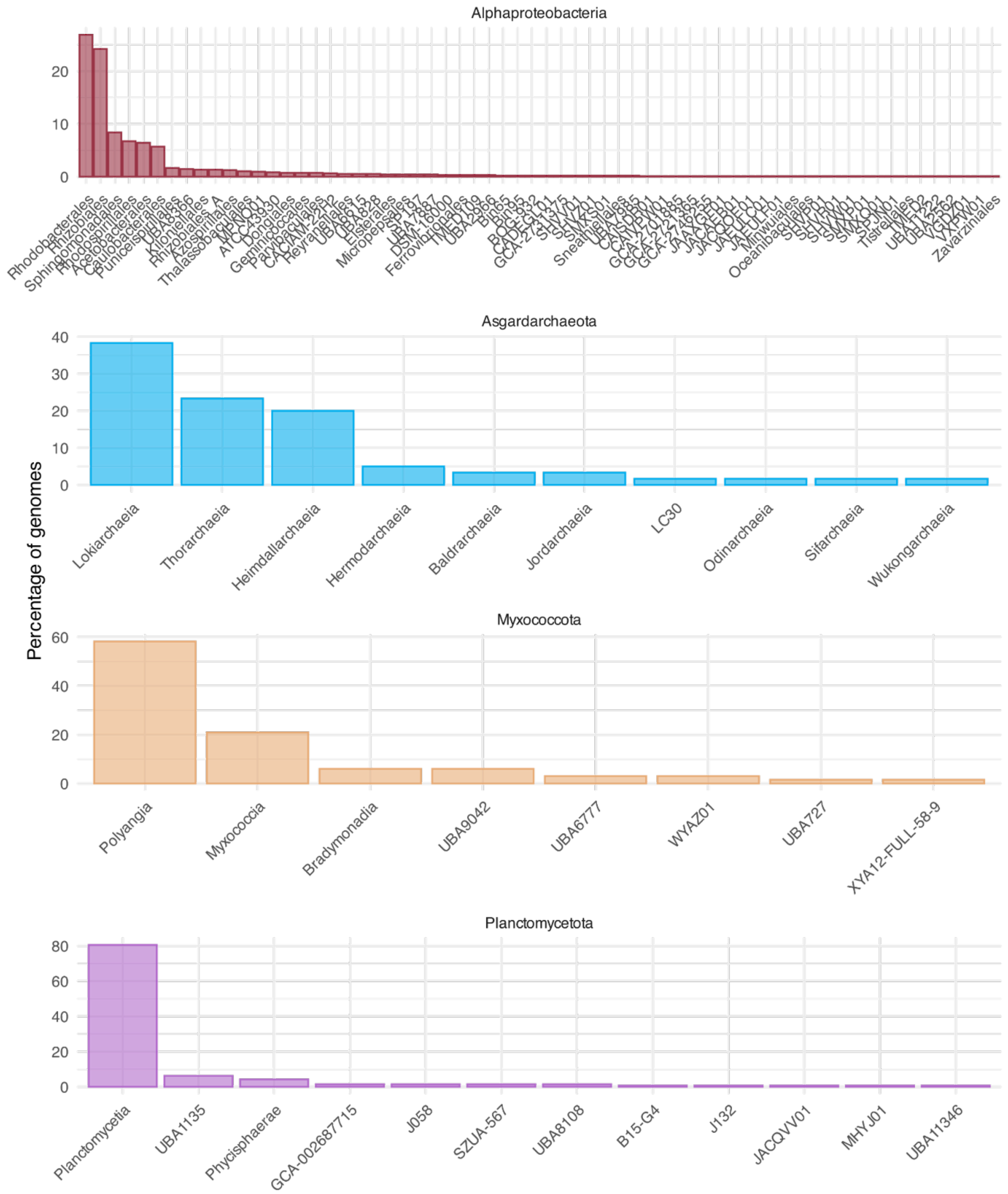

**Supplementary Fig. 12. Taxonomic distribution of extant genomes of donor groups employed for the inference of metabolic traits**. Percentage of genomes per class- (for Asgardarchaeota, Planctomycetota, Myxococcota) or order- (for Alphaproteobacteria) level lineage.

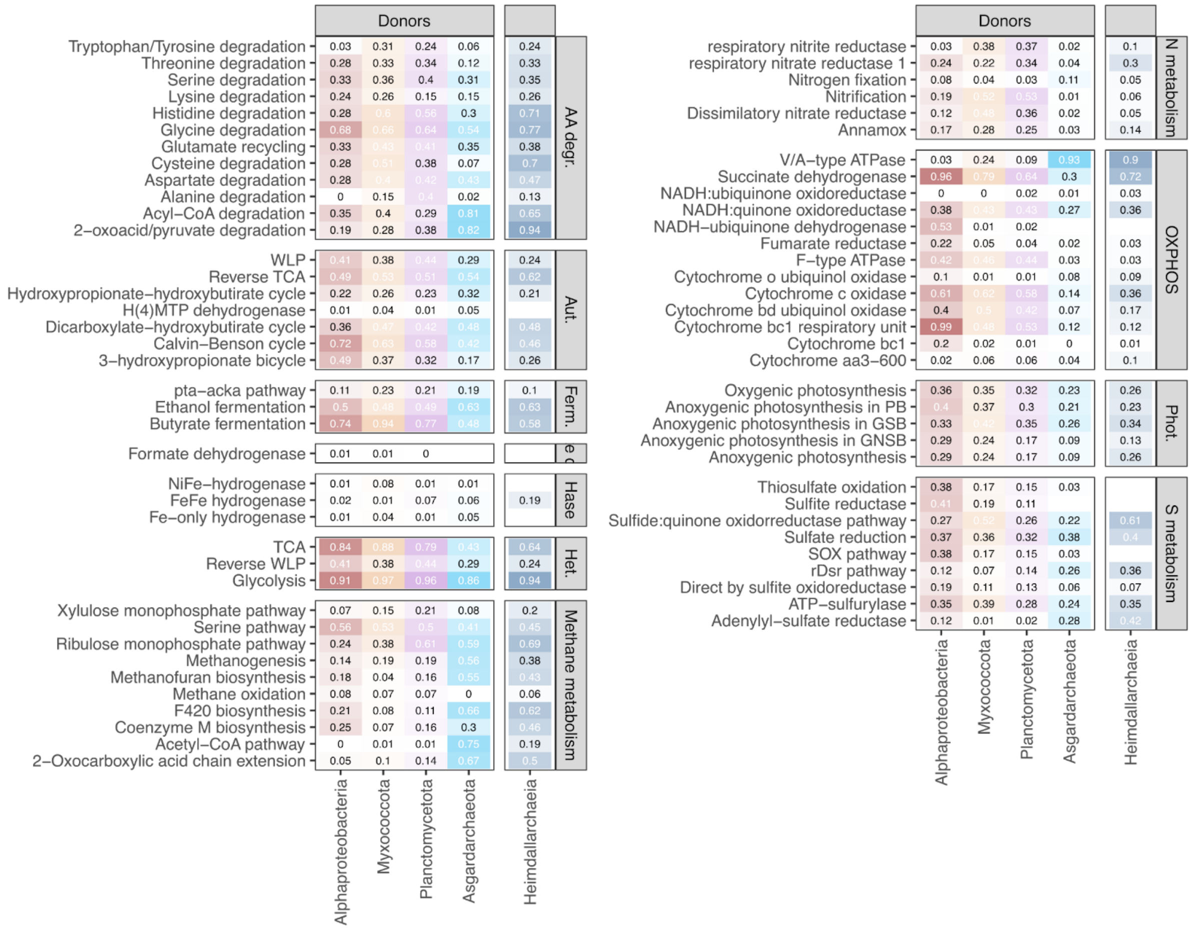

**Supplementary Fig. 13: Prevalence of the pathways and enzymes in the genomes of the donor's descendants for the resampling approach**. Each box shows a group of metabolic pathways and enzymes. The opacity of the colour shows the prevalence (the percentage of genomes with the feature) of that specific pathway or enzyme in extant genomes from the donor's clades that share at least 50% of the KOs that these donors transferred to LECA. Abbreviations, AA degr.: amino acid degradation; Aut.: autotrophy, Ferm.: fermentation; Hase: hydrogenases; Het.: heterotrophy; N: nitrogen metabolism; OXPHOS: oxidative phosphorylation; Phot.: photosynthesis; S metabolism: sulphur metabolism.

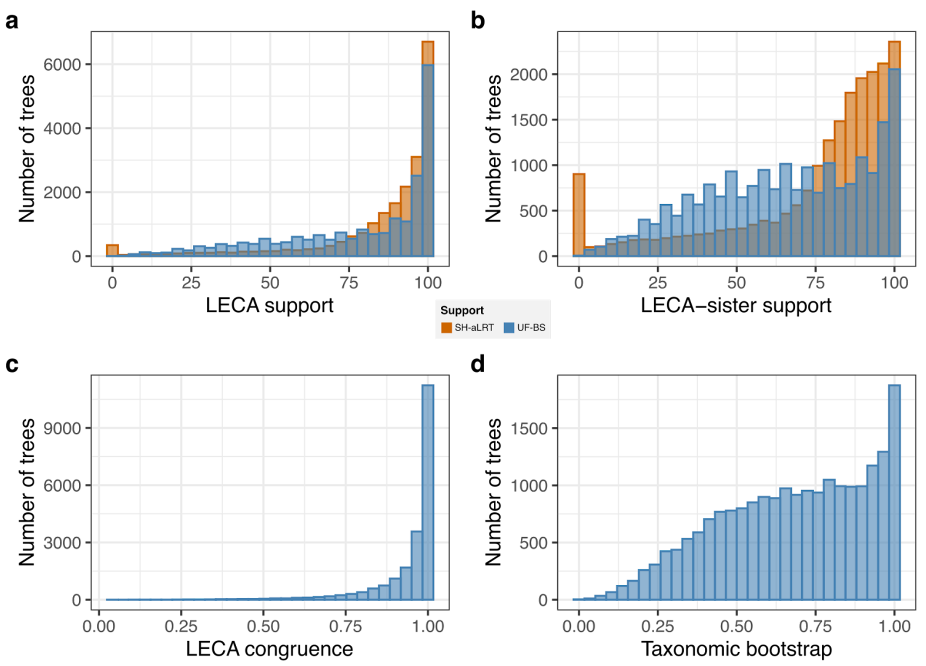

**Supplementary Fig. 14. Summary of the variables considered for the "stress test".** Barplots representing the number of trees included in different measures: a) Support for the branch subtending the LECA clade, columns in blue represent UltrafastBS and columns in orange represent SH-aLRT. b) Support values for the branch subtending the clade formed by the LECA clade and its sister clade, columns in blue represent UltrafastBS and columns in orange represent SH-aLRT c) the mean Jaccard distance of the LECA groups in the bootstrap trees and the maximum likelihood (ML) tree LECA group, a distance of 0 would mean that both LECA definitions do not share any sequence and 1 they are identical. d) The proportion of bootstrap trees that agree with the ML donor.

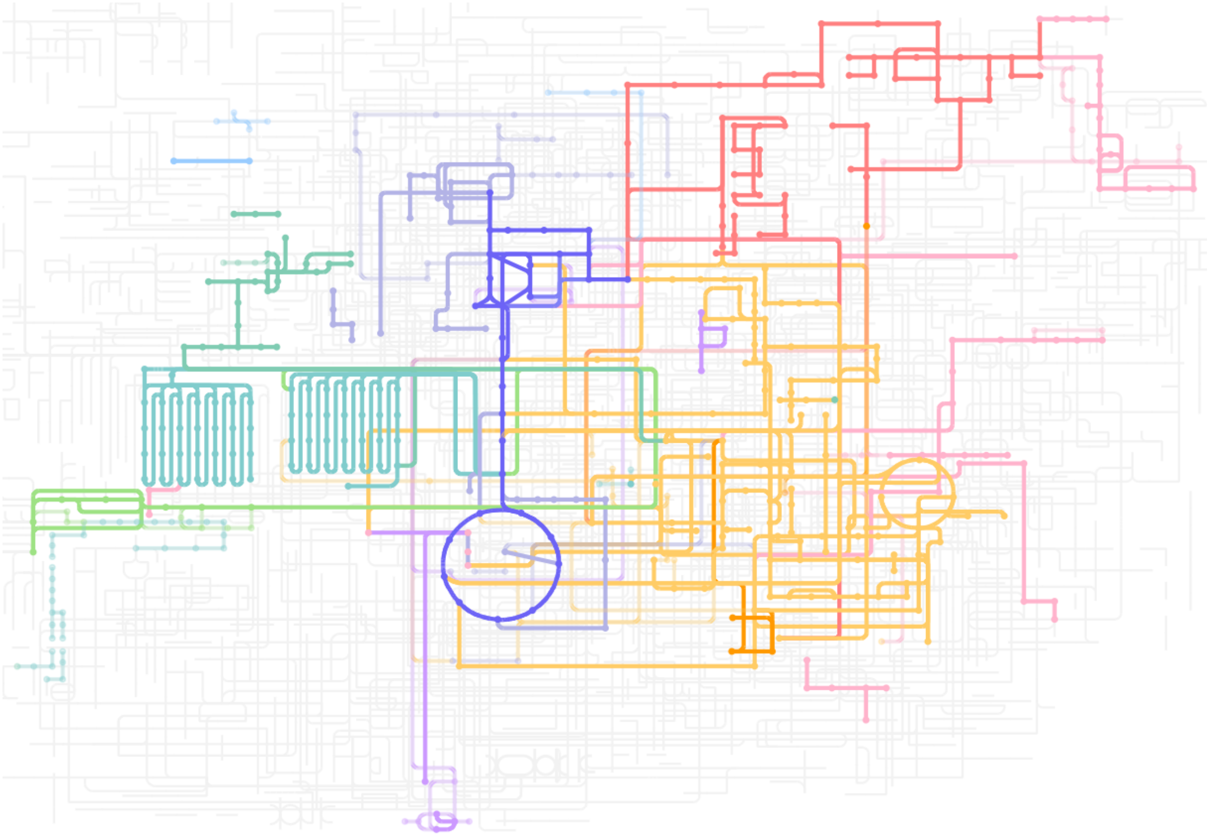

**Supplementary Fig. 15. Impact of taxonomy-aware automatic and manual filtering of terms on the inferred LECA metabolism.** The full colour set of metabolic modules inferred from the LECA metaproteome filtered by taxonomic breadth (at least two eukaryotic kingdoms, according to KEGG) and manual curation (we excluded three modules involved in carbon fixation and included V-type ATPase, whose KEGG definition just includes animals). In lighter colour, additional complete metabolic modules in LECA directly inferred from the metaproteome.

#### Supplementary Tables

**Supplementary Table 1**. Genomes considered in each eTOLDB database with their sources, the supergroup and division considered in this study, the filtered proteome size and the lineage.

**Supplementary Table 2**. Summary of the taxonomic groups considered in the study and the number of genomes in each before and after filtering and those incorporated in eTOLDB.

**Supplementary Table 3**. Summary of the number of LECA gene families in the Orthogroup and mLECA-OG steps of the pipeline and their origins for the latter.

**Supplementary Table 4**. Inferred metabolic modules present in LECA according to the reconstructed consensus annotated proteome (KOs inferred in LECA). Complete and incomplete pathways are shown, as well as the most recent common ancestor (MRCA) calculated using the KEGG taxonomy. Those modules specific of narrow eukaryotic groups or specific of prokaryotic particular functions were flagged to have not been present in LECA (column: In LECA).

**Supplementary Table 5**. Data on shared number of KOs between LECA metaproteome and the eukaryotic genome of species within the Free living unicellular osmotrophs (FLUO), autotrophs (FLUA) and phagotrophs (FLUP). COG frequencies for each proteome and COG functional category are also included.

**Supplementary Table 6**. List of innovations. Innovations are those KOs annotated in trees of any database and for which we do not find any non-eukaryotic homolog. They are further restricted to those KOs that are pan-eukaryotic according to KEGG.

**Supplementary Table 7**. Proportion of the consensus proteome KOs from a certain origin that are involved in the different features.

**Supplementary Table 8.** List of acquired donors through virus mediated HGT.

#### Supplementary Data availability

All the alignment, trees, annotations, and supporting files to carry on this research are deposited in a Zenodo repository that will be available upon publication.
